# Macrophage-secreted Pyrimidine Metabolites Confer Chemotherapy Resistance in Acute Myeloid Leukemia (AML)

**DOI:** 10.1101/2025.11.01.686055

**Authors:** Chuqi Wang, Yuhan Wang, Camillo Benetti, Xiao Xian Lin, Enes Dasdemir, Petra Hyroššová, Jordan Yong Ming Tan, Edward Ayoub, Karanpreet Singh Bhatia, Fang Qi Lim, Yun Li, Zihui Zhao, Ahmed M. Mamdouh, Shir Ying Lee, Jakub Rohlena, Chad V. Pecot, Michael Andreeff, Katerina Rohlenova, Hussein A. Abbas, Shruti Bhatt

## Abstract

The tumor microenvironment (TME) programs cancer cells to influence therapeutic responses. Macrophages residing in TME switch from pro-phagocytic to tumor-promoting and immunosuppressive phenotypes as cancer develops. While these pro-tumor functions of macrophages are associated with poor outcomes, the underlying mechanisms by which bone-marrow (BM)-associated macrophages fuel myeloid malignancy and their precise contribution to relapse remain undissected. Here, we show expansion of monocyte/macrophage population in leukemia patients post-chemotherapy relapse, and spatial proximity of macrophages to leukemia blasts in the BM niche. This proximity proved functionally consequential—depletion of macrophages delayed leukemia relapse post cytarabine (AraC), a frontline chemotherapy, in patient-derived xenografts (PDX) and syngeneic leukemia models. Mechanistically, a pyrimidine metabolite, deoxycytidine (dC), secreted by BM macrophages, is taken up by leukemia cells to directly inhibit deoxycytidine kinase (DCK) to hamper AraC activation and subsequent resistance in a cell non-autonomous manner. Diagnosis AML patients exhibited significantly higher circulating dC levels than healthy donors, and dC levels further increased following chemotherapy. *SAMHD1,* which catalyzes deoxynucleoside triphosphates (dNTPs) into deoxynucleoside, was highly abundant in macrophages and mediated dC accumulation. Blockade of dC production in mouse and human macrophages via genetic and pharmacological inhibition of *SAMHD1* or *DHODH*, a critical enzyme in pyrimidine synthesis, restored AraC sensitivity. Combination with DHODH inhibitors significantly delayed AraC relapse in human PDX and mouse syngeneic AML models. Collectively, we identify a metabolic immune–leukemia crosstalk in which *SAMHD1*^high^ macrophages mediates chemoresistance by secreting pyrimidine metabolites and propose macrophage metabolic reprogramming as a tractable strategy to overcome TME-driven chemoresistance in myeloid leukemia.

## Introduction

Acute myeloid leukemia (AML) is an aggressive hematologic malignancy characterized by the uncontrolled expansion of immature myeloid blasts spanning a spectrum of differentiation states, from early hematopoietic stem and progenitor cells to terminally differentiated populations [1]. For over four decades, the “7+3” chemotherapy regimen, a combination of cytarabine (AraC) and an anthracycline (typically daunorubicin), has demonstrated remarkable success in achieving initial remission, yet its long-term efficacy remains limited. More than 50% of patients who achieve remission during initial treatment experience relapse within months to years after treatment, highlighting the persistent challenge of durable disease control [2, 3].

While most studies addressing chemoresistance in AML have focused on genomic aberrations, recent evidence suggests that the immunosuppressive bone marrow microenvironment (BME) plays a pivotal role in cancer therapy resistance. Among BME components, macrophages are of particular interest as these innate immune cells can acquire tumor-supportive phenotypes in many cancers [4–6]. However, the role of macrophages in AML remains muddier, particularly regarding whether macrophages contribute to or attenuate the therapeutic efficacy of anti-AML therapies. Several studies suggest that macrophages play a pro-tumor role in AML [7, 8]. Furthermore, M2-like macrophages are associated with poor prognosis [8, 9]. Depletion of CD169+ macrophages in a mouse AML model enhanced the response to chemotherapy [10]. Conversely, other studies suggest macrophages retain anti-leukemic activity through phagocytosis. Depletion of macrophages using clodronate liposomes in immunodeficient and immunocompetent mice led to an increase in AML burden [11–13], while mice lacking LC3-associated phagocytosis (LAP) had a faster AML expansion [11]. Taken together, the existing literature suggests that macrophages adapt to diverse functions in the BM niche; however, their precise role, underlying mechanisms, and contribution to relapse development remain underdetermined.

Here, we reveal that macrophages were enriched in AML patients at relapse and are spatially proximal in BME. Integration of genome-wide CRISPR screen in AML cells and global metabolomics identified that macrophages secrete deoxycytidine (dC), a pyrimidine metabolite to competitively inhibit deoxycytidine kinase (DCK) in AML cells, thereby blocking cytarabine activation (Ara-CTP) and initiating resistance. Single-cell sequencing of healthy and AML samples, and cancer-wide Dependency Map (DepMap) analysis identified that *SAMHD1,* that catalyzes dNTP degradation, is highly abundant in macrophages and causing dC accumulation. Inhibition of dihydroorotate dehydrogenase (DHODH) can suppress macrophage-derived dC production and restore AraC sensitivity in vivo, providing a potential therapeutic strategy. Overall, this finding presents a critical contribution of macrophages in AML chemoresistance and provides novel opportunity to develop treatment strategies that selectively target the metabolic phenotype of bone marrow macrophages.

## Results

### Macrophages are associated with chemoresistance and counteract AraC efficacy in vivo

To identify key cell types that contribute to AML poor outcomes, we analyzed scRNA-seq, bulk RNA-seq, and spatial transcriptomics data from primary AML bone marrow (Figure 1A). To identify microenvironmental cell types associated with chemotherapy relapse, we analyzed scRNA-seq data from paired AML bone marrow (BM) samples collected at diagnosis and relapse after AraC-based therapy (Figure 1B–C, Table S1, S1A–C). Monocytes/macrophages expanded significantly in relapsed samples (Figure 1C and S1A). To test cell types associated with survival, we examined bulk RNA-seq data from the BeatAML cohort. Focusing on samples with <30% blasts to minimize leukemia cells’ influence, we correlated immune gene signatures with patient survival (Figure S2). High scores for M0/M2 macrophages and dendritic cells, but not M1 macrophages, correlated with shorter survival, whereas T-and B-cell signatures correlated with longer survival (Figure S2) [14]. Using an independent published macrophage signature [4], we again found that higher macrophage score was associated with poor outcome (Figure 1D).

**Figure 1.**
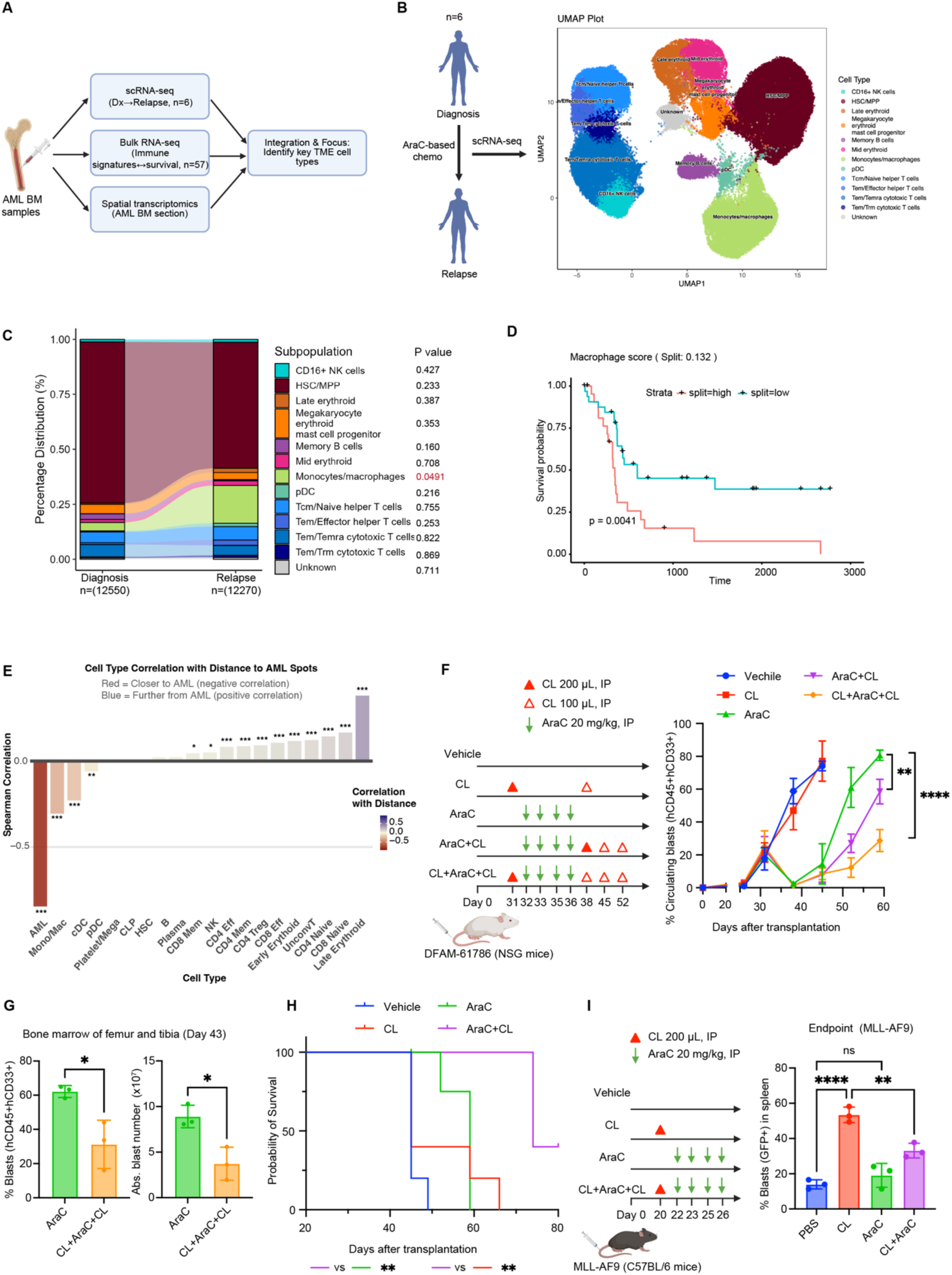
Macrophages counteract AraC efficacy in vivo. (A) Schematic figure of bioinformatic analyses that use scRNA-seq, bulk RNA-seq, and spatial transcriptomic data to identify key TME cell types. (B) Schematic of sample collection for scRNA-seq and UMAP visualization of bone marrow cells at diagnosis vs. relapse. (C) Proportion of major cell types at diagnosis and relapse. Paired t test. (D) Kaplan–Meier survival curves of AML patients (BeatAML; ≤30% blasts in BM; n = 57) stratified by macrophage gene signature. Log-rank test. (E) Correlation between cell type abundance and distance to AML regions. Negative correlation indicates closer localization to AML cells. (F) Macrophage depletion in the PDX model DFAM-61786 using clodronate liposomes (CL). Leukemia engraftment over time in DFAM-61786 PDX mice (n = 5 per group). Unpaired t test. (G) Percentage and absolute number of hCD45+hCD33+ PDX cells in bone marrow after treatment (n = 3). Unpaired t test. (H) Kaplan–Meier survival curves of DFAM-61786 PDX mice after treatment. (I) Macrophage depletion in MLL-AF9 syngeneic AML using CL. Percentage of GFP+ blasts in spleens of treated mice (n = 3 per group). One-way ANOVA. *Data are mean ± SD. *, p < 0.05; **, p < 0.01; ***, p < 0.001; ****, p < 0.0001.

To investigate the spatial interaction between AML blasts and stromal cells, we analyzed the spatial transcriptomics of an AML BM section published in a previous paper (Figure S3A) [15]. We stratified the BM section into 5 zones according to distance to AML cells, core_AML, adjacent, proximal, intermediate, and distant (Figure S3A-B). We observed that monocytes/macrophages (Mono/Mac) were preferentially localized adjacent to AML-rich areas, while T and NK cells were spatially excluded (Figure 1E, S3C-G). To confirm the annotation of these Mono/Mac, we calculated the expression of macrophage score using an independent gene signature and found high expression of macrophage genes in Mono/Mac but not AML cells (Figure S3H). Together, these data suggest close interaction between AML cells and macrophages that may influence chemotherapy response.

To test the functional role of macrophages, we depleted them with clodronate liposomes (CL) in an AML patient-derived xenograft (PDX) model, DFAM-61786 (Figure 1F, S4A-B). Macrophage depletion significantly reduced tumor burden in AraC-treated mice but not vehicle-treated mice, suggesting that macrophages cause AraC resistance but do not influence leukemia growth (Figure 1F-G). Macrophage depletion also prolonged the survival of AraC-treated mice (Figure 1H). In vitro, CL had no direct toxicity and did not affect AML cell viability (Figure S4C). In contrast to AraC, CL did not alter response to the BCL2 inhibitor Venetoclax (Ven) in vivo, indicating drug-specific resistance caused by macrophages (Figure S4D–E). To investigate the effect of macrophages in immunocompetent setting, we depleted macrophages in mice bearing MLL-AF9 syngeneic AML (Figure 1I). Since MLL-AF9 cells are refractory to AraC, low-dose AraC did not reduce tumor burden in macrophage-competent mice, however, AraC can reduce burden in CL-treated mice (Figure 1I and S4F), again suggesting that macrophages mediate AraC resistance. Surprisingly, macrophage depletion alone significantly increased tumor burden, implying that macrophages also exert anti-tumor activity in vivo, likely via immune activation. Together, these findings demonstrate that macrophages specifically drive resistance to AraC, though their depletion is not therapeutically viable due to loss of anti-tumor functions and toxicity.

### Macrophages release small-molecule metabolites to inhibit AraC activation

To assess whether macrophages directly protect AML cells from AraC, we cocultured AML cells (MOLM13, PDX, and primary AML cells) with mouse bone marrow–derived macrophages (BMDM). To differentiate the effects of soluble factors and direct cell-cell contact, we cocultured macrophages and AML cells by direct coculture or transwell coculture (Figure 2A). Both direct and transwell coculture increased AML survival under AraC, indicating contributions from soluble factors as well as direct cell–cell contact (Figure 2B). Without AraC, transwell coculture had minimal effect on increasing AML cell number (Figure S5A). Coculture with human monocyte-derived macrophages (hMDM) increased PDX cell survival under DMSO or Ven treatment, but to a greater extent under AraC treatment (Figure 2C).

**Figure 2.**
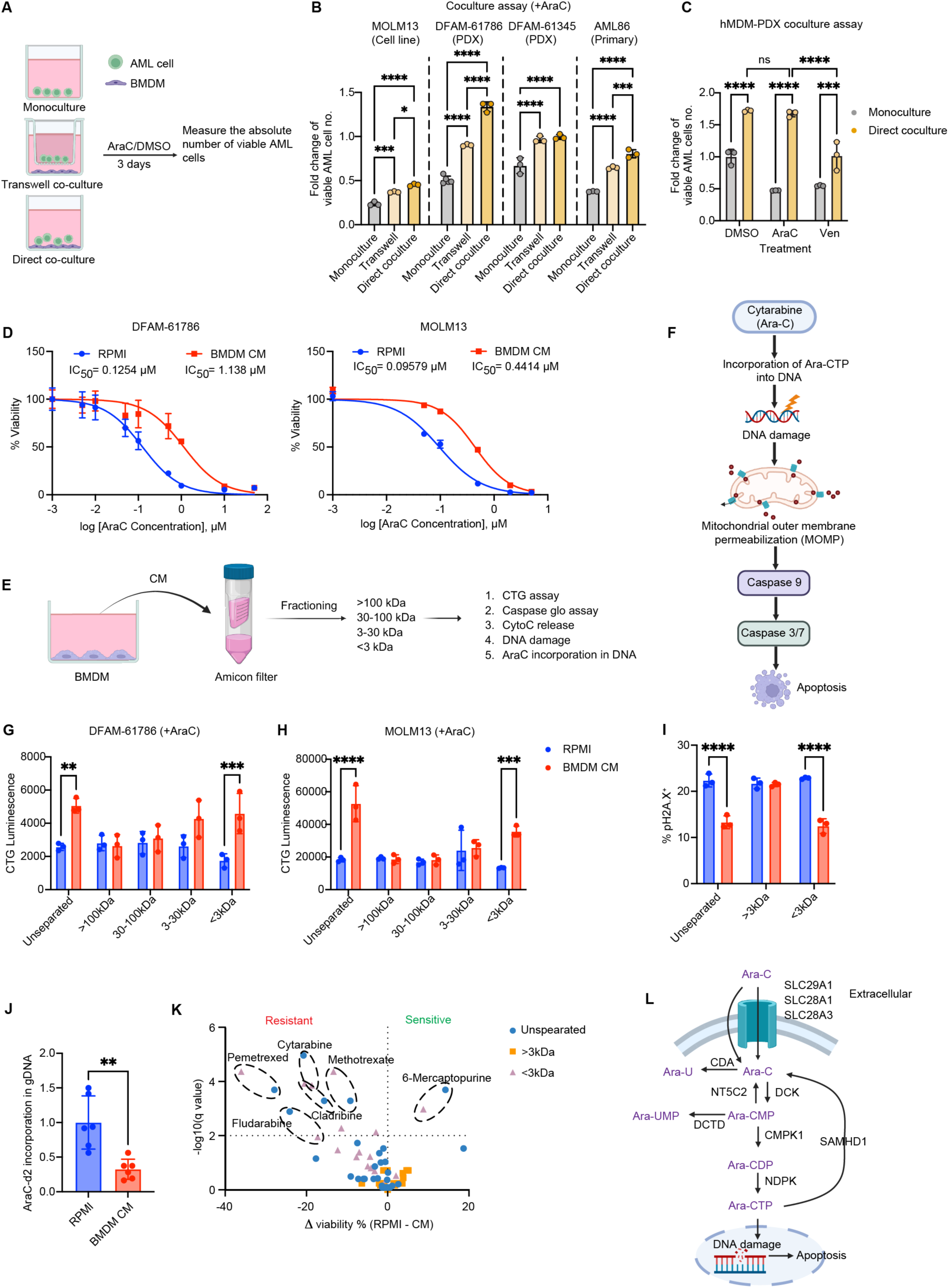
Macrophages release small-molecule metabolites that inhibit AraC activity. (A) Schematic of macrophage-AML coculture assays with direct contact or transwell separation. (B) Absolute number of live MOLM13 cells in monoculture vs. coculture under AraC. One-way ANOVA. (C) Fold change of absolute number of live DFAM-61786 PDX cells in monoculture vs. coculture with human monocyte-derived macrophages (hMDM) under DMSO, AraC, or Ven treatment. (D) Viability of DFAM-61786 or MOLM13 cells treated with AraC ± 50% BMDM CM. (E) Schematic of CM fractionation by ultrafiltration. BMDM CM or RPMI control media were fractioned using Amicon® Ultra Centrifugal Filters to obtain fractions with different molecular weights. (F) Pathway schematic of AraC-induced DNA damage and apoptosis. (G and H) Using CellTiter-Glo® (CTG) assay for the quantification of viable DFAM-61786 or MOLM13 cells under AraC with BMDM CM or RPMI fractions. (I) ãH2AX staining after AraC ± CM fractions. Two-way ANOVA. (J) Incorporation of AraC-d2 into genomic DNA in MOLM13 cells with CM vs. RPMI, quantified by LC-MS. Unpaired t test. (K) Drug screening with fractions of BMDM CM or RPMI using MOLM13 cells. (L) Schematic figure showing the metabolism pathway of AraC in cancer cells before incorporation into genomic DNA. *Data are mean ± SD. *, p < 0.05; **, p < 0.01; ***, p < 0.001; ****, p < 0.0001.

To validate that soluble factors from macrophages can mediate AraC resistance, we supplemented conditioned media (CM) from macrophages to AML culture. CM from BMDM (BMDM CM) reproduced the protective effect in AraC alone and AraC+daunorubicin combination treatment (Figure 2D, S5B). To investigate what components can cause resistance, we fractioned the CM of BMDM and RPMI control media using ultrafiltration (Figure 2E). Surprisingly, although macrophages can secrete various cytokines and chemokines including osteopontin (Figure S5C), only the <3kDa fraction, primarily small-molecule metabolites, of BMDM CM conferred AraC resistance (Figure 2G-H). Notably, neither boiling nor nuclease treatment abolished the protective activity of BMDM CM, suggesting that the soluble factors are heat-stable and not nucleic acids (Figure S5D-E).

AraC is a pyrimidine analog that can incorporate into genomic DNA, finally leading to DNA damage and cell death [16–18]. To investigate at which level do the soluble factors block AraC-mediated apoptosis, we analyzed different activities in the intrinsic apoptosis pathway (Figure 2F). Mechanistically, the <3 kDa CM fraction of CM attenuated AraC-induced caspase-3/7 and caspase-9 activation, cytochrome C (CytoC) release, and γH2AX DNA damage (Figure 2I, S5F-G). Furthermore, caspase3/7 inhibition was comparable between direct and transwell coculture under AraC, indicating that soluble metabolites—not cell-cell contact—block apoptosis (Figure S5I). CM also reduced incorporation of isotope-labeled AraC into genomic DNA (Figure 2J), indicating blockade of AraC activation upstream of DNA incorporation by CM.

To test whether soluble factors of macrophages can cause resistance to other drugs, we conducted a drug screening with 32 drugs in the presence of fractions of CM or RPMI control. The drug screening revealed that the <3kDa fraction also reduced sensitivity to other antimetabolites (methotrexate, pemetrexed, fludarabine, and cladribine), while sensitizing to 6-mercaptopurine, suggesting a potential mechanism linked to nucleotide metabolism (Figure 2K).

AraC needs to be phosphorylated by several kinases into the active form Ara-CTP before incorporation into DNA (Figure 3G). Disruption of AraC metabolism into the Ara-CTP form can reduce AraC efficacy [19–22]. Notably, RNA-seq of AML cells after coculture or in vivo macrophage depletion showed no significant transcriptional changes in AraC-metabolizing genes (Figure S6A-E), indicating that macrophage-derived metabolites may regulate AraC metabolism at post-translational levels.

**Figure 3.**
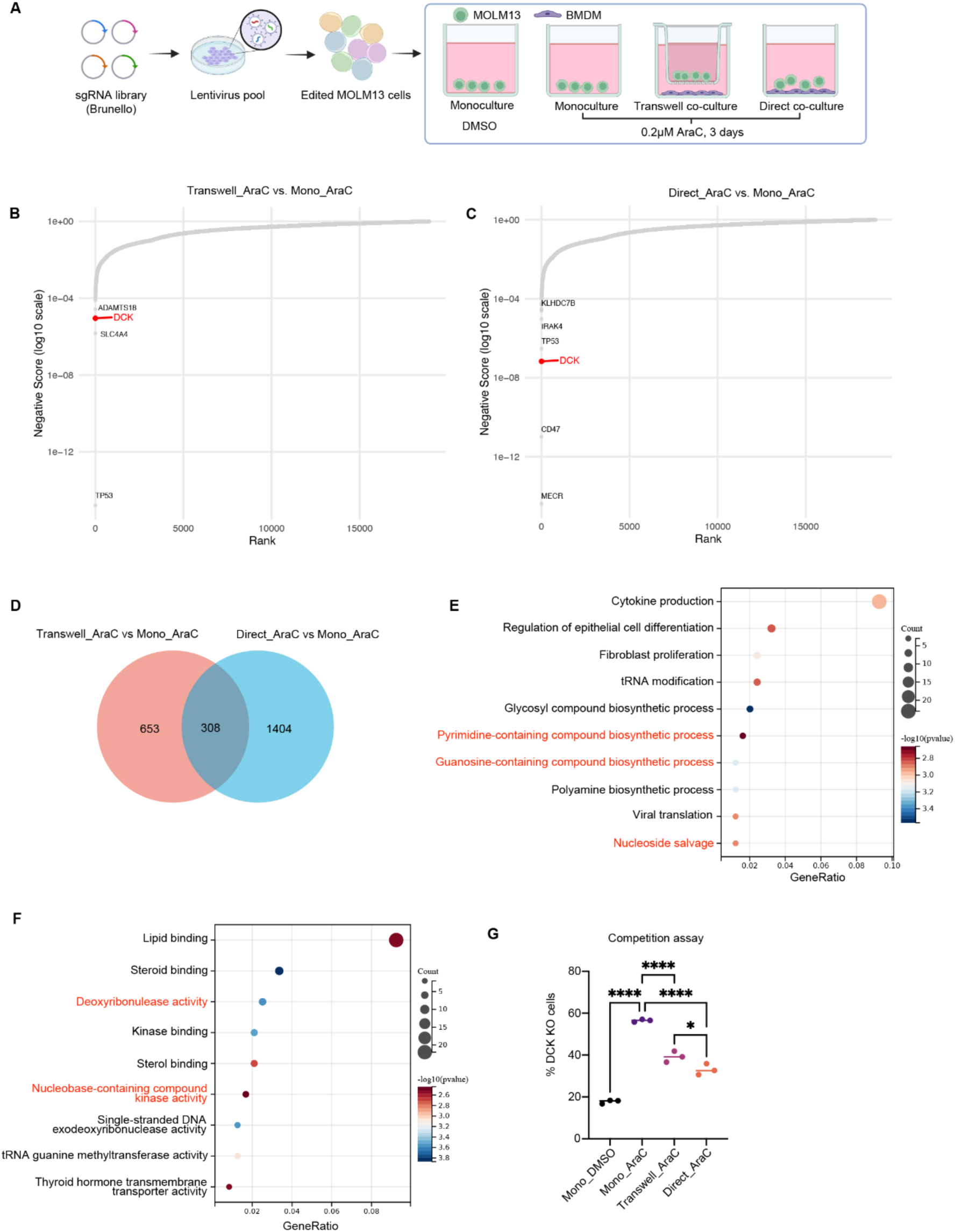
Genome-wide CRISPR screen identifies DCK as the target of macrophage-derived metabolites. (A) Schematic of CRISPR screen in MOLM13 cells with macrophage coculture. (B and C) Scatter plots showing sgRNAs depleted in Transwell_A vs. Mono_A and in Direct_A vs. Mono_A. (D) Venn diagram showing shared depleted top hits (p<0.05) between Transwell_A vs. Mono_A and in Direct_A vs. Mono_A. (E) Gene Ontology (GO) term enrichment analysis of common hits in D. (F) KEGG pathway enrichment of common hits in D. (G) The % of DCK KO cells in the competition assay.

### Intercellular CRISPR screen identifies DCK as the node of influence by macrophage-derived metabolites

To identify the molecular target of macrophage-derived metabolites, we performed a genome-wide CRISPR screen in MOLM13 cells (Figure 3A). As expected, comparison of AraC-treated monoculture (Mono_AraC) vs. DMSO-treated monoculture (Mono_DMSO) recapitulated known AraC resistance/sensitizing genes (e.g., SAMHD1, BCL2, TP53, BAX) and pathways (e.g., Cell cycle, DNA replication) [23], confirming the robustness of the screen (Figure S7A-C).

When comparing AraC-treated transwell coculture (Transwell_AraC) with Mono_AraC and AraC-treated direct coculture AraC (Direct_AraC) with Mono_AraC, we identified common sgRNA hits in both comparisons which are potential target genes of macrophage-derived metabolites (Figure 3B-D). Gene set enrichment analysis of common significant hits highlighted an enrichment in pyrimidine and nucleoside metabolism (Figure 3E-F, S7D). Interestingly, DCK was identified as a common top hit associated with pyrimidine and nucleoside metabolism (Figure 3B-C). It is a critical enzyme that phosphorylates AraC into its active form Ara-CTP (Figure 2L) [24]. Loss of DCK is a well-established mechanism of AraC resistance (Figure S7E) [23, 25]. To validate the finding from the CRIPSR screen, we performed a competition assay using DCK knockout (KO) cells mixed with empty vector (EV) cells (Figure S7F). As expected, DCK KO cells expanded in Mono_AraC compared to Mono_DMSO, but this enrichment was diminished in Transwell_AraC and Direct_AraC (Figure 3G), mirroring the CRISPR screen results. These data suggest that while loss of DCK protects AML cells from AraC, the protective effect is reduced in macrophage cocultures because macrophage-derived metabolites already inhibit DCK activity. Collectively, these results indicate that macrophages may release metabolites to antagonize DCK function, thereby impairing AraC activation.

### Macrophages secrete the pyrimidine metabolite deoxycytidine (dC) to block AraC activation and efficacy

To identify macrophage-released metabolites responsible for AraC resistance, we performed untargeted metabolomics and found deoxycytidine (dC) to be one of the most enriched metabolites in BMDM CM compared to RPMI control (Figure 4A–B). Analysis of the DepMap database further revealed a strong positive correlation between intracellular dC abundance and AraC IC50 across cancer cell lines (Figure 4C, S8A), implicating dC as a key metabolite associated with AraC resistance. Thus, we focused on dC which is both enriched in macrophage CM and closely linked to AraC resistance (Figure 4D). Targeted measurement by HPLC/MS confirmed the presence of dC in macrophage CM and mouse BM aspirates but not in control media and AML CM (Figure 4E). Supplementing AML cultures with exogenous dC at levels equivalent to those detected in CM reproduced macrophage-mediated AraC resistance (Figure 4F). This result indicates that dC is the primary metabolite in BMDM CM that blocks AraC activity. Consistent with our findings, it has also been shown by others that dC can block DCK activity and cause resistance to other antimetabolite drugs [26, 27]. Newly diagnosed AML patients exhibited significantly higher blood dC levels than healthy donors, and dC levels further increased following AraC treatment (Figure 4G), suggesting the clinical relevance of this finding. Importantly, physiological concentrations of dC observed in patient blood were sufficient to induce AraC resistance in both MOLM13 and primary AML cells (Figure 4H–I). Exogenous dC also induced resistance to fludarabine, but had no effect on pemetrexed, methotrexate, or 6-mercaptopurine (Figure S8B–E).

**Figure 4.**
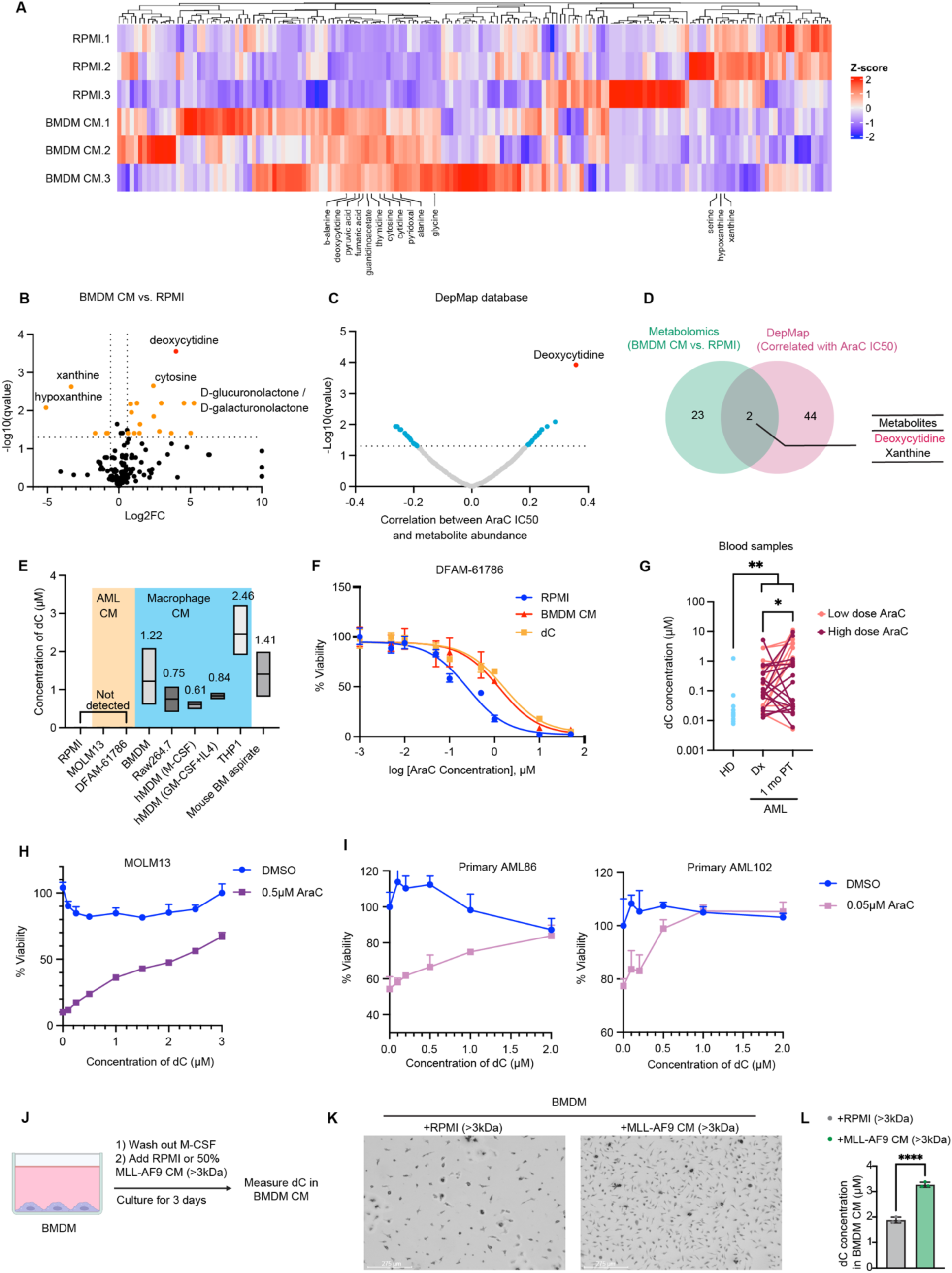
Macrophages secrete pyrimidine metabolite deoxycytidine (dC) that blocks AraC activation and efficacy. (A-B) Heatmap and volcano plot of metabolites enriched in BMDM CM vs. RPMI. (C) Correlation of metabolite abundance with AraC IC50 in cancer cell lines (DepMap). Deoxycytidine showed strongest correlation with AraC IC50. Pearson correlation. (D) Venn diagram illustrating overlap between metabolites significantly altered in BMDM CM (metabolomics) and those significantly correlated with AraC IC50 (DepMap). (E) Targeted measurement of dC concentrations in RPMI control media, CM from AML cells (MOLM13, DFAM-61786), CM from macrophages (BMDM, Raw264.7, hMDM, THP1), and mouse BM aspirates. (F) Viability of DFAM-61786 AML cells under AraC with either RPMI control, BMDM CM, exogenous dC. (G) Targeted measurement of dC levels in blood samples from healthy donors (HD), newly diagnosed (Dx), and one-month post-treatment (1 mo PT) AML patients. (H) Viability of MOLM13 treated with AraC ± exogenous dC at physiological concentrations. (I) Viability of two primary AML samples treated with AraC ± exogenous dC at physiological concentrations. (J) Schematic figure showing the experimental design of treating BMDM with CM from MLL-AF9 leukemic cells or RPMI control media. The CM from BMDM were collected after treatment for dC measurement. (K) Representative microscopic images showing the BMDM confluency after M-CSF withdrawal and treatment with RPMI or MLL-AF9 CM (>3kDa). (L) Increased dC release by BMDM after treatment with MLL-AF9 CM (>3kDa). *Data are mean ± SD. *, p < 0.05; **, p < 0.01; ***, p < 0.001; ****, p < 0.0001.

To test whether AML cells can influence the amount of dC released by macrophages, we supplemented CM from MLL-AF9 leukemic cells into BMDM culture after M-CSF withdrawal (Figure 4J). The large molecules (>3kDa) in CM from MLL-AF9 AML cells maintained BMDM survival in the absence of M-CSF (Figure 4K) and increased dC release (Figure 4L), suggesting that AML cells can “educate” macrophages to produce more dC.

### SAMHD1 drives macrophage-derived dC production

To determine whether macrophages are the main source of dC in vivo, we depleted macrophages with CL (Figure 5A). This significantly reduced dC concentrations in bone marrow aspirates, confirming macrophages as the principal source (Figure 5B). Analysis of scRNA-seq datasets (van Galen et al.) revealed that SAMHD1, a dNTP hydrolase that converts dNTPs into deoxynucleosides, is highly expressed in proMono, Mono/Mac and dendritic cells (Figure 5C–E, S9A–D), consistent with previous reports of high SAMHD1 expression in myeloid populations [28–30]. DepMap correlation analysis further identified SAMHD1 as one of the top genes correlated with intracellular dC abundance (Figure 5F–G). We therefore hypothesize that SAMHD1 can drive the biosynthesis of dC in macrophages.

**Figure 5.**
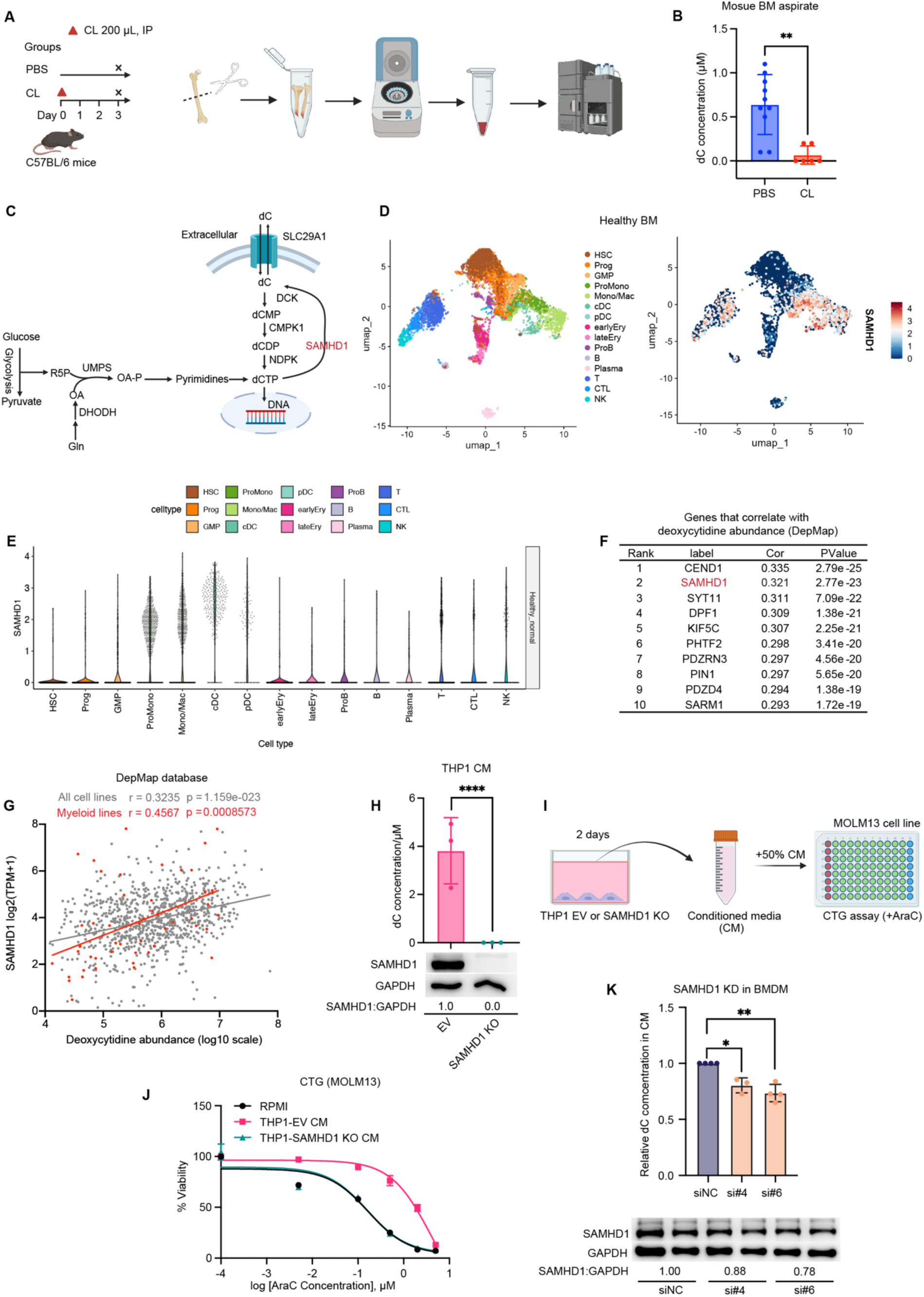
SAMHD1 drives macrophage-derived dC production. (A) Schematic of in vivo experiment to test macrophage contribution to dC. (B) dC concentration in BM aspirates of mice treated with PBS vs. CL (PBS n=10, CL n=6). (C) Schematic of dC biosynthesis pathway. (D and E) scRNA-seq analysis of healthy BM showing high SAMHD1 expression in myeloid cells (ProMono, Mono/Mac, DC). (F) Top 10 genes that correlate with dC abundance across cancer cell lines (DepMap). SAMHD1 is among top hits. (G) DepMap correlation of SAMHD1 expression with dC abundance across all cancer cell lines (grey) or myeloid lines (red). Pearson correlation. (H) SAMHD1 KO in THP1 human monocyte/macrophage cell line abolished dC release. Paired t test. (I) Schematic figure showing the experimental design of CTG using CM from THP1 WT vs. SAMHD1 KO. (J) Viability of MOLM13 cells under AraC with CM from THP1 EV vs. SAMHD1 KO. (K) siRNA-mediated SAMHD1 knockdown in BMDM reduced dC secretion. One-way ANOVA. *Data are mean ± SD. *, p < 0.05; **, p < 0.01; ***, p < 0.001; ****, p < 0.0001.

SAMHD1 KO in THP1 human monocyte/macrophage cell line completely abolished dC secretion and reversed CM-mediated AraC resistance (Figure 5H–J, S9F–H). Knocking down of SAMHD1 in BMDM using siRNA also reduced dC release (Figure 5K and S9I). These data suggest that SAMHD1 is essential to produce dC by macrophages. Notably, SAMHD1 is also expressed in differentiated AML subpopulations resembling monocytes/macrophages and dendritic cells, suggesting that mature myeloid-like malignant cells themselves may contribute to dC secretion (S9B and S9D).

### SAMHD1 is upregulated in relapse and induced by inflammation

To test whether SAMHD1 upregulation is associated with chemotherapy relapse, we analyzed SAMHD1 expression in paired primary AML samples (Figure 1B). scRNA-seq analysis revealed increased SAMHD1 expression in Mono/Mac after relapse (Figure 6A–B). Spatial transcriptomics showed high SAMHD1 expression in AML-rich and adjacent zones, suggesting that leukemia cells may be exposed to locally elevated dC (Figure 6C, S3A, S9E). Consistently, a prior study reported that SAMHD1^high^ Mono/Mac in newly diagnosed AML BM predicted early relapse in pediatric AML [31].

**Figure 6.**
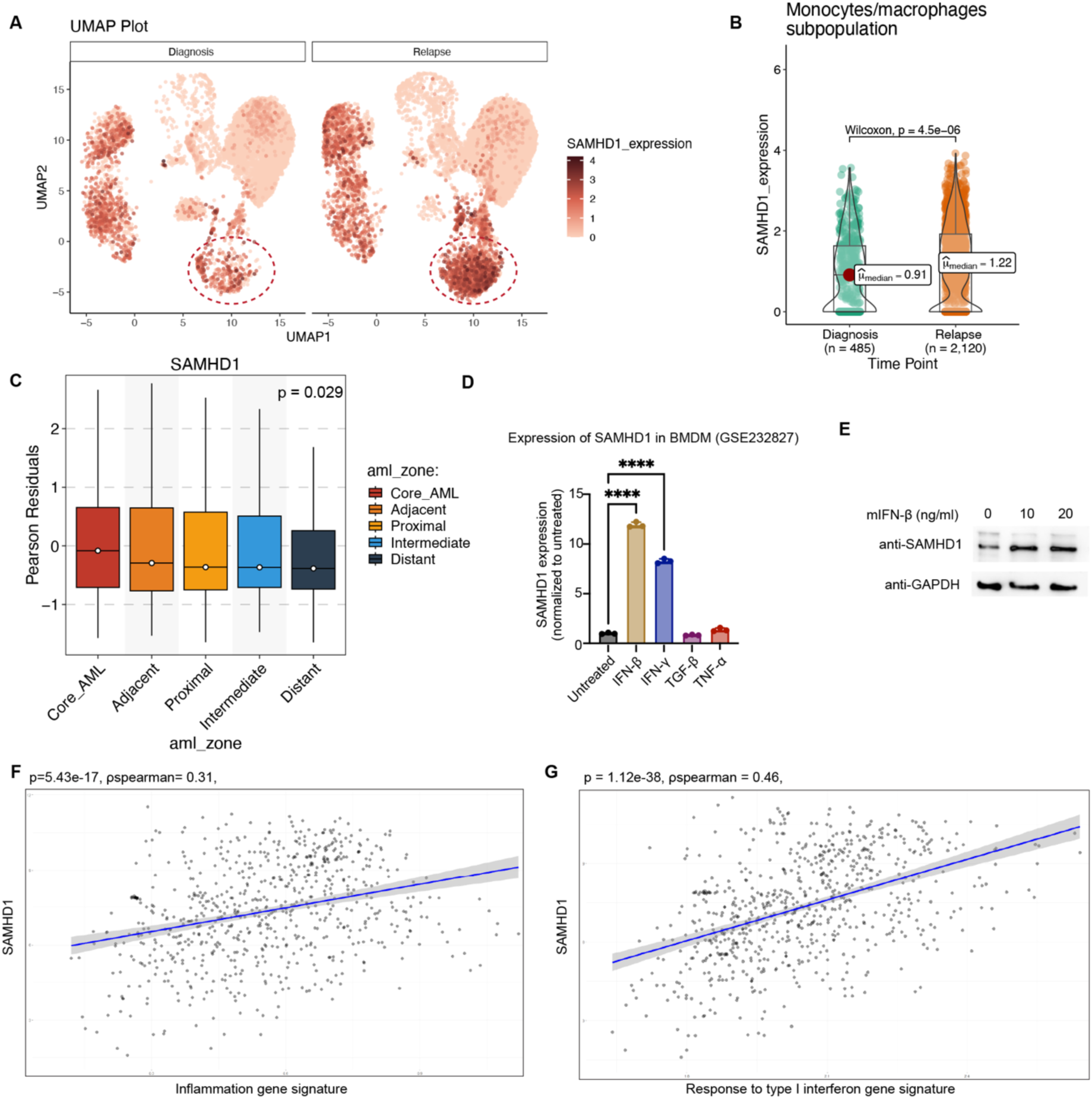
SAMHD1 is upregulated after relapse and induced by inflammation. (A) UMAP of paired AML BM samples showing expansion of SAMHD1^high^ Mono/Mac after relapse. (B) scRNA-seq quantification of SAMHD1 expression in Mono/Mac cells at diagnosis vs. relapse. (C) Spatial transcriptomics showing high SAMHD1 expression in AML-rich core zones in BM. (D) Expression of SAMHD1 in BMDM treated with IFN-β, IFN-γ, TNF-α, or TGF-β (GSE232827). Only interferons upregulated SAMHD1. (E) Western blot confirming SAMHD1 induction by IFN-β in BMDMs. (F and G) Correlation of SAMHD1 expression with inflammation and type I IFN gene signatures in BeatAML dataset. *Data are mean ± SD. *, p < 0.05; **, p < 0.01; ***, p < 0.001; ****, p < 0.0001.

Previous studies have shown that inflammation is associated with poorer overall survival in AML [32]. Moreover, interferon-gama signaling is associated with and is a predictive biomarker for chemoresistance [33]. However, the mechanisms by which inflammation and interferon signaling affecting chemotherapy efficacy are not known. Because SAMHD1 is an interferon-stimulated gene [34], we investigated whether inflammatory signaling regulates SAMHD1 expression. Indeed, type I and II interferons upregulated SAMHD1 in BMDM, whereas TNF-α and TGF-β had no effect (Figure 6D–E). In BeatAML bulk RNA-seq dataset, SAMHD1 expression also correlated strongly with inflammation and interferon-response signatures (Figure 6F–G). These data suggest that inflammation regulates the expression of SAMHD1 in macrophages and is a potential mechanism of chemoresistance in AML.

### DHODH inhibition reduces dC production and restores AraC sensitivity

After elucidating the mechanism, we next sought to overcome the resistance by inhibiting dC production in the microenvironment. Given the lack of SAMHD1 inhibitors with good draggability, we targeted upstream pyrimidine biosynthesis by inhibiting DHODH, a critical enzyme in the pyrimidine biosynthesis pathway (Figure 5C). DHODH inhibitor Leflunomide has been approved for the treatment of rheumatoid arthritis [35], while more DHODH inhibitors are being tested for the treatment of cancer [36]. DHODH inhibition has been shown to differentiate AML in pre-clinical studies [37, 38], however, clinical data of DHODH inhibitors showed futility as a monotherapy in treating AML and other cancers [36, 39–41].

To test whether DHODH inhibition can reduce the secretion of dC by macrophages, we measured dC in CM from BMDM induced from WT or DHOHD-/-mouse BM. Genetic deletion in BMDM significantly reduced dC release (Figure 7A). Similarly, DHODH KO in THP1 cell line significantly reduced dC release and reversed CM-mediated AraC resistance (Figure 7B-D). In addition to genetic disruption, pharmacologic inhibition with Leflunomide (Lef) or Brequinar (BRQ) similarly reduced dC in CM and restored AraC sensitivity in vitro (Figure 7E-G).

**Figure 7.**
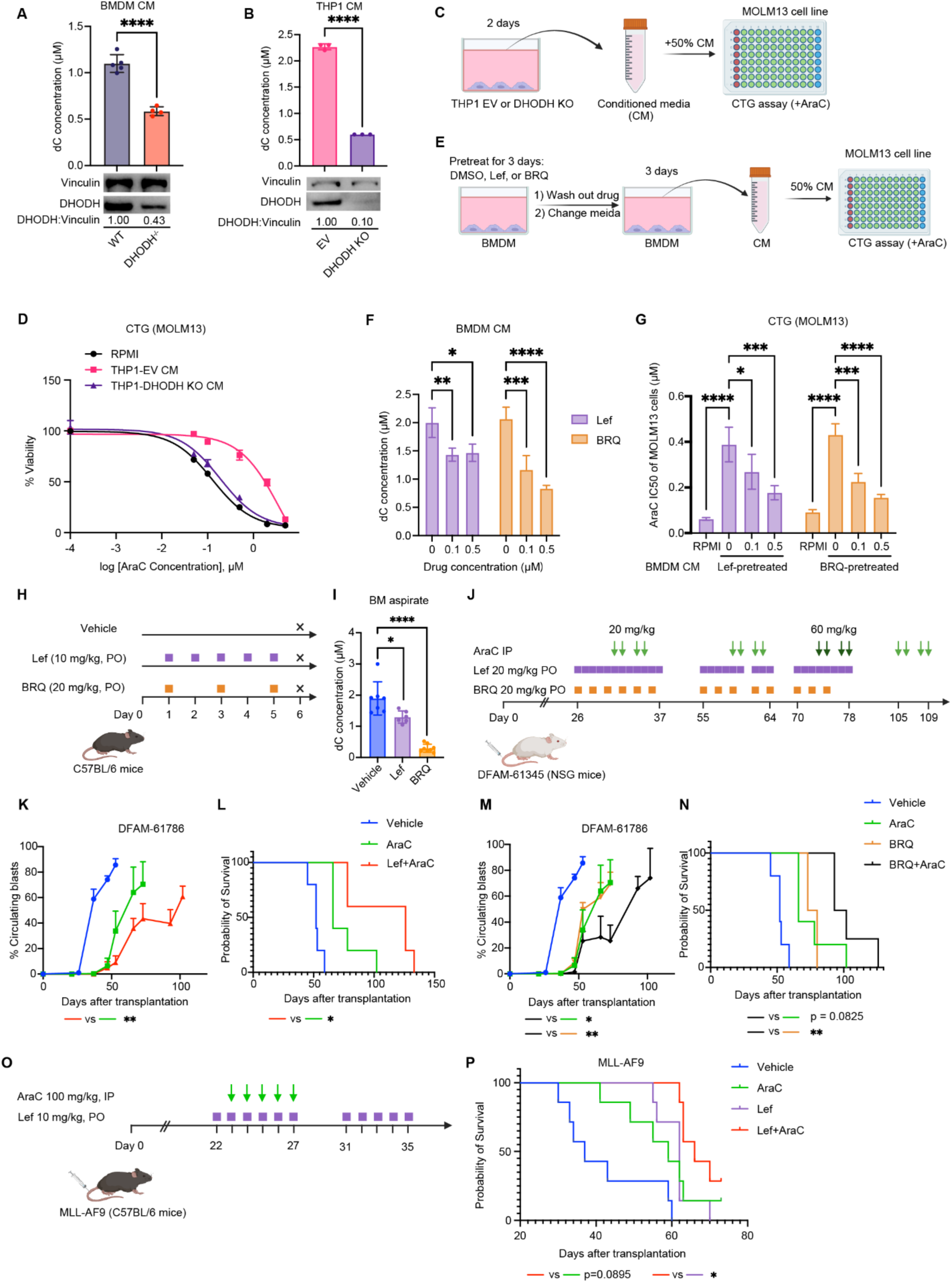
DHODH inhibition enhances the efficacy of AraC. (A) Genetic deletion of DHODH in BMDMs reduced dC release. (B) DHODH KO in THP1 cells similarly reduced dC secretion. (C-D) MOLM13 viability under AraC with CM from THP1 EV vs. DHODH KO. (E) Design of in vitro assay testing the effect of DHODH inhibitors on macrophage dC release and AraC response. (F) dC concentrations in BMDM CM after pretreatment with Lef or BRQ. (G) AraC IC50 of MOLM13 cells with CM from BMDM pretreated with Lef or BRQ. (H and I) DHODH inhibitors lowered dC levels in mouse BM aspirates. (J) Schematic of DFAM-61786 PDX model testing AraC + DHODH inhibitors. (K-N) Leukemia engraftment and survival curves of DFAM-61786 PDX mice under AraC vs. AraC + DHODH inhibitors. (O-P) Survival of MLL-AF9 mice treated with AraC vs. AraC + Leflunomide. Data are mean ± SEM in K and M. Data in other figures are mean ± SD. *Data are mean ± SD. *, p < 0.05; **, p < 0.01; ***, p < 0.001; ****, p < 0.0001.

In vivo, both inhibitors lowered dC levels in bone marrow aspirates (Figure 7H–I). Combining AraC with Lef or BRQ significantly reduced leukemia burden and prolonged survival in the PDX model DFAM-61786 (Figure 7J – 7N). Combining Lef also prolonged the survival in the AraC-refractory MLL-AF9 model (Figure 7O-P). Since Leflunomide monotherapy showed little efficacy in prior trials, we omitted single-agent groups to minimize animal use. Together, these findings demonstrate that DHODH inhibition suppresses macrophage-derived dC and enhances the therapeutic activity of AraC.

## Discussion

The TME is increasingly recognized as a driver of drug resistance in hematologic malignancies, although its role has long been underestimated [42–44]. In AML, macrophages have been associated with poor outcomes [7–10, 45], but the mechanisms behind this link have remained unclear. Here, we identify a metabolic macrophage–leukemia axis in which macrophages secrete the pyrimidine metabolite dC to block AraC activation (Figure 8). This mechanism of chemoresistance is likely not restricted to AraC but may extend to other nucleoside analogs that depend on DCK, such as fludarabine, decitabine and cladribine [24, 46]. Beyond dC, our metabolomic profiling also revealed alterations in other metabolites in macrophage CM, including depletion of hypoxanthine, which could explain the altered response to 6-mercaptopurine [47]. These findings suggest that macrophages influence a broader spectrum of drug sensitivities through nucleoside/nucleotide metabolic interactions.

**Figure 8.**
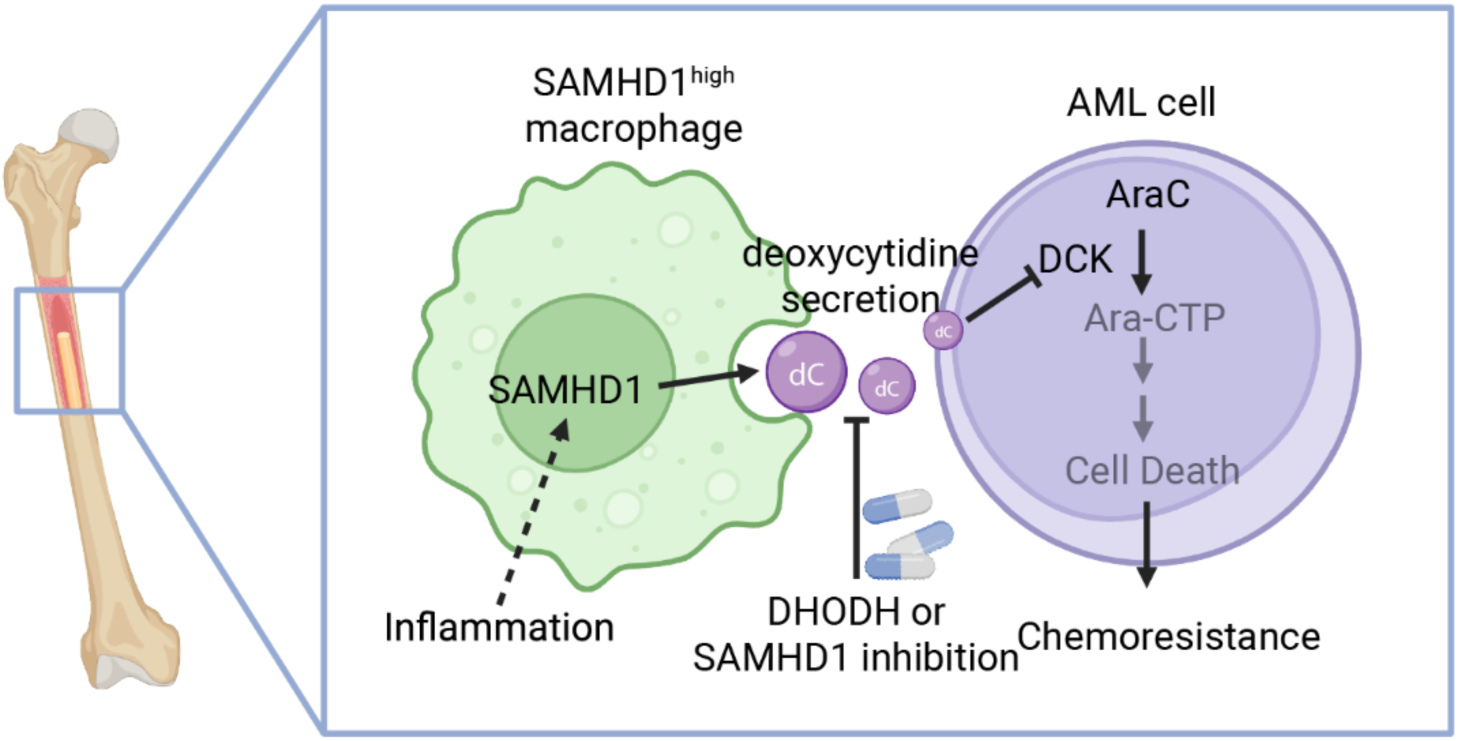
Schematic figure of metabolic crosstalk between BM macrophages and AML cells in promoting AraC chemoresistance. SAMHD1^high^ macrophages secrete dC to block DCK in AML cells, and thereby, inhibit the activation of AraC and subsequent cell death. Inflammation, specifically IFN, can upregulate SAMHD1 expression in macrophages and is a potential mechanism that underlies chemoresistance. Inhibition of DHODH or SAMHD1 in macrophages could decrease microenvironmental dC supply and sensitize AraC.

Interestingly, a prior study in pancreatic ductal adenocarcinoma (PDAC) showed that macrophages can release dC and promote resistance to gemcitabine, a standard-of-care chemotherapy in PDAC [27]. However, in blood malignancies, it is still unknown whether macrophages are the dominant in vivo source of bone marrow environmental dC and why macrophages produce dC. Our findings reveal that dC secretion is driven by high expression of SAMHD1, a dNTP hydrolase known as an antiviral restriction factor in myeloid cells [28, 29, 48]. Thus, dC production may be a byproduct of the innate immune system’s defense mechanisms, exploited by leukemia to evade chemotherapy.

Prior studies have shown that AML-intrinsic SAMHD1 directly drives AraC resistance by hydrolyzing Ara-CTP [21, 22]. Our findings extend the AraC resistance mechanism of SAMHD1 by showing that tumor-extrinsic SAMHD1 in macrophages also promotes AraC resistance, indirectly through production of extracellular dC. Our scRNA-seq analysis further revealed that differentiated AML subpopulations with myeloid-like features also express high levels of SAMHD1, suggesting that subpopulations of AML cells themselves may contribute to dC secretion.

Inflammation has a paradoxical role in cancer. Chronic inflammatory signaling can drive tumorigenesis and therapy resistance. Previous studies have shown that inflammatory signatures in AML correlate with poor survival and reduced chemotherapy efficacy [32, 33, 49]. Macrophages are a major source of inflammatory transcripts in AML bone marrow, further implicating them in this process. We confirmed that both type I and type II interferons upregulated SAMHD1 expression in macrophages. This provides a mechanistic explanation for how inflammation contributes to chemoresistance. Consistently, a previous study showed that interferons induce SAMHD1 and elevate dC abundance in cells [50].

From a therapeutic perspective, we showed that simply depleting macrophages may impair the good functions of macrophages in immunocompetent mice and cause toxicity. While SAMHD1-specific inhibitors are lacking, we therefore proposed an alternative strategy by targeting upstream pyrimidine biosynthesis through DHODH inhibition. Genetic and pharmacological inhibition of DHODH reduced dC secretion by macrophages and restored AraC sensitivity in vitro and in vivo. Importantly, DHODH inhibitors such as leflunomide and brequinar are already clinically approved or are under investigation in clinical trials, with well-established safety and pharmacokinetic profiles. While DHODH inhibitors have shown limited efficacy as monotherapies in AML, our data suggest that their use in combination with AraC could be effective by simultaneously lowering dC-mediated resistance.

## Methods

See supplemental information.

## Acknowledgements

S.B. acknowledges support from the Singapore Ministry of Education (MOE) Academic Research Fund (AcRF) Tier 2 (MOE-000573-00), NUS-Paris University Joint Grant, the Department of Pharmacy and Pharmaceutical Sciences Starup Funds at National University of Singapore. S.B. is a recipient of the EMBO Global Investigator Award.

We thank Xiaoning Wang and Delia Pang at the MD1 Flow Cytometry core facility of the National University of Singapore for assistance with instruments and cell sorting. We thank Peng Gao at the Metabolomics core facility, Feinberg School of Medicine, Northwestern University for assistance with metabolomics. We thank Lin Tan at the Metabolomics core facility of MD Anderson for assistance with analyzing patient samples. Jennifer Marvin-Peek and Jessica Root for preparing patient blood samples. Adam Sperling for providing the MLL-AF9 cells.

## Author contributions

C.W. conceptualized the study, designed the experiments, performed experiments, analyzed data, and wrote the manuscript. Y.W. designed the experiments, performed experiments, analyzed data, and wrote the manuscript. S.B. conceptualized the study, designed the experiments, and wrote the manuscript. C.B., J.Y.M.T., P.H., F.Q.L., Y.L., and A.M.M. performed experiments. X.X.L., E.D., E.A., K.S.B, and Z.Z. analyzed sequencing experiments and plotted data. S.Y.L. provided patient samples. M.A. provided scRNA-seq data of paired AML samples. A.A.H provided patient samples and spatial transcriptomics data. K.R., J.R., and P.H. provided DHODH-/-mice and designed the experiments.

