## Supplemental Figures 1-8, Methods and Materials for "Macrophage-secreted Pyrimidine Metabolites Confer Chemotherapy Resistance in Acute Myeloid Leukemia (AML)"

### Supplemental information

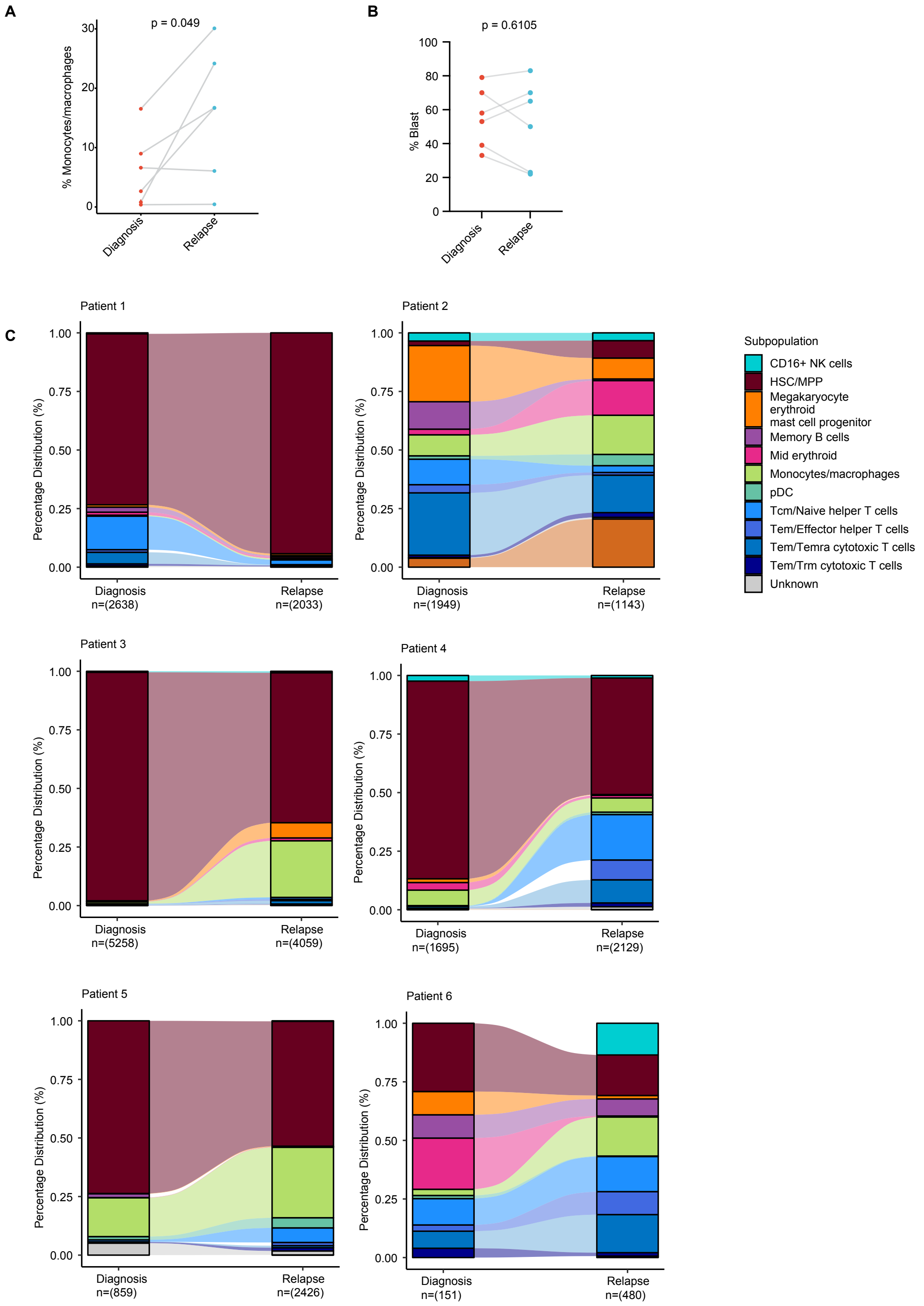

Supplemental figure 1. Single-cell RNA sequencing (scRNA-seq) analysis of paired primary AML bone marrow samples.

(A) Percentage of Mono/Mac in paired AML BM samples at diagnosis vs. relapse (n = 6). Paired t test.

(B) Percentage of blasts in paired AML BM at diagnosis vs. relapse (n = 6). No significant difference was observed, indicating that changes in Mono/Mac are not due to altered disease burden. Paired t test.

(C) Proportion of cell types at diagnosis and relapse from 6 patients.

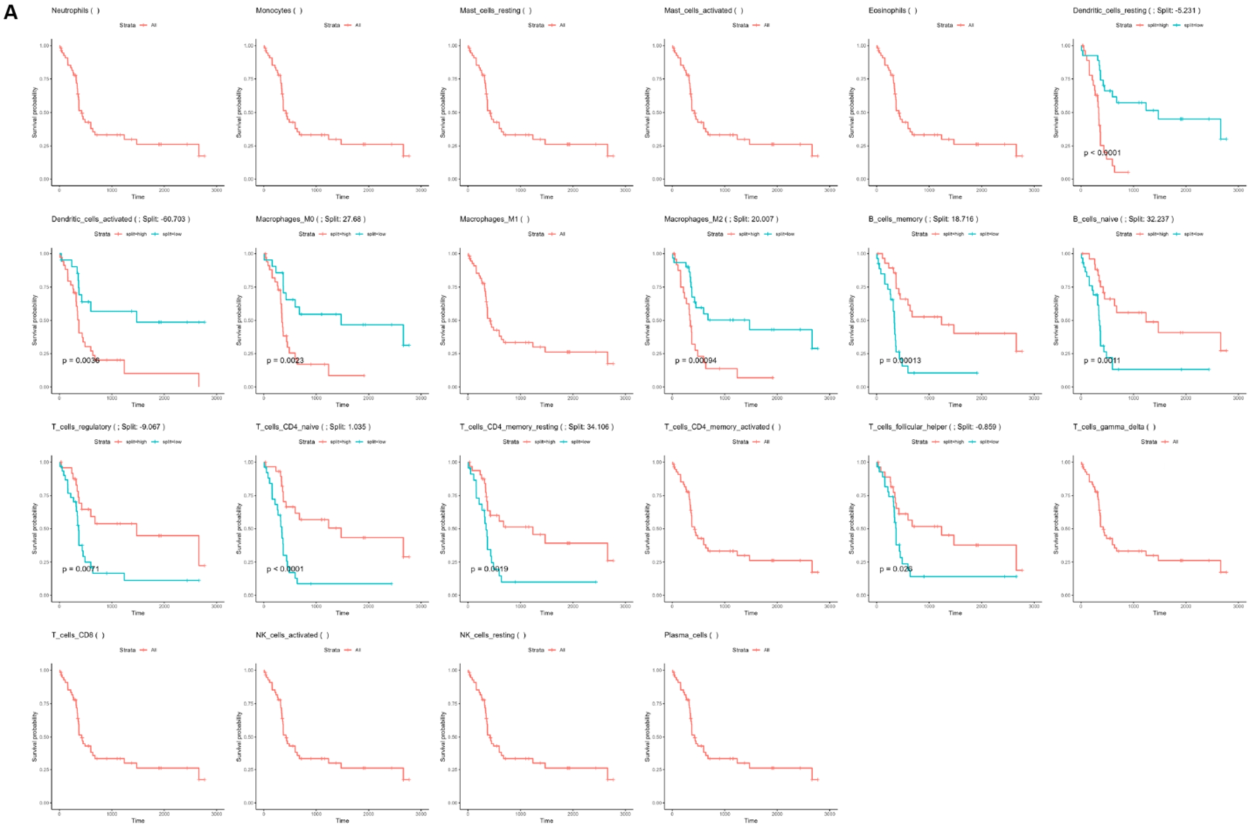

Supplemental figure 2. Kaplan–Meier curves of AML patients (BeatAML cohort; ≤30% BM blasts; n = 57) stratified into high vs. low immune cell populations using LM22 gene signature and random forest classification. One curve suggests no significant difference in overall survival between high vs. low groups.

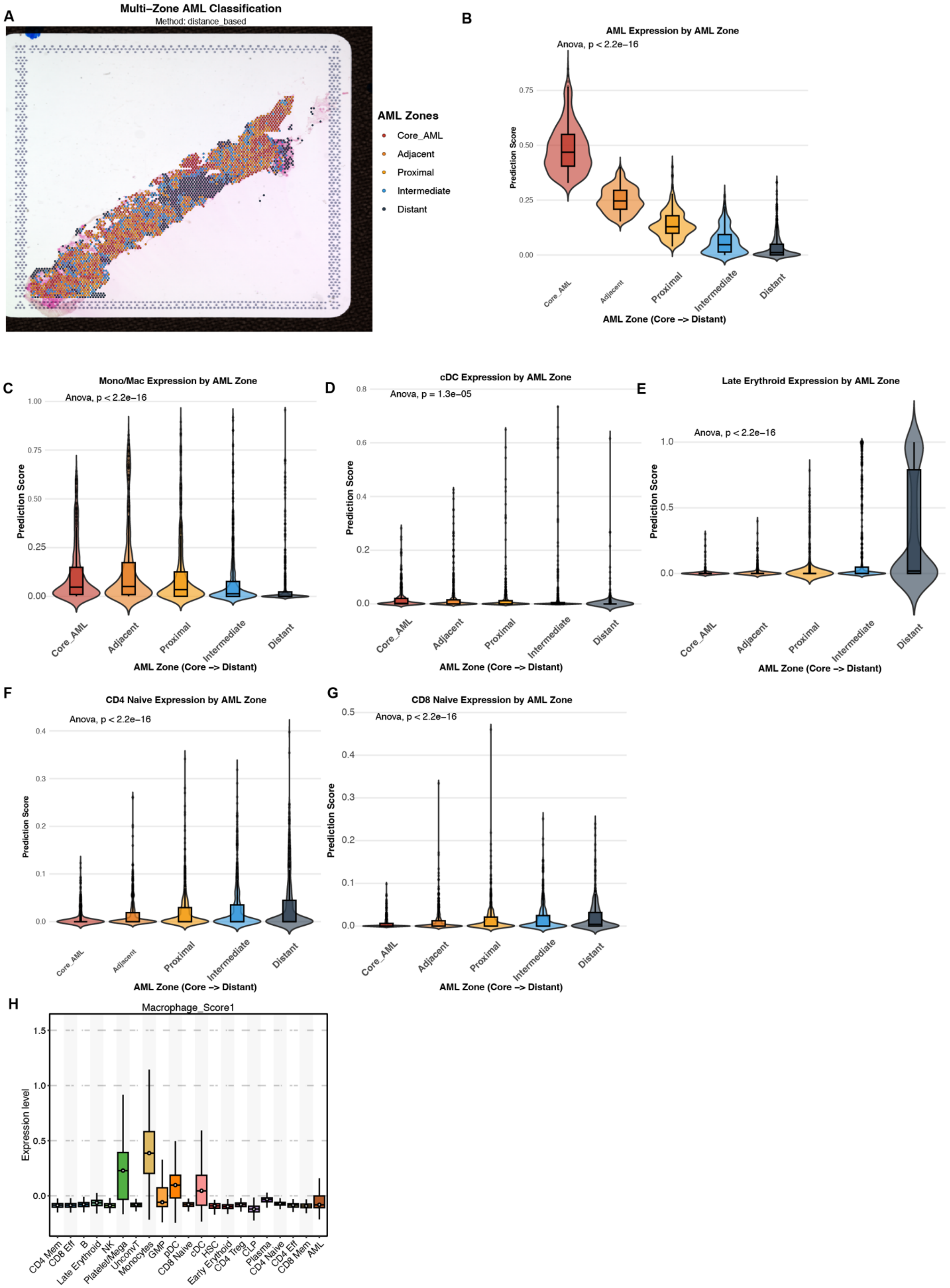

A
Multi−Zone AML Classification
Method: distance_based
AML Zones
Core_AML
Adjacent
Proximal
Intermediate
Distant
B
AML Expression by AML Zone
Anova, p < 2.2e−16
Prediction Score
AML Zone (Core −> Distant)
C
Mono/Mac Expression by AML Zone
Anova, p < 2.2e−16
D
cDC Expression by AML Zone
Anova, p = 1.3e−05
E
Late Erythroid Expression by AML Zone
Anova, p < 2.2e−16
F
CD4 Naive Expression by AML Zone
Anova, p < 2.2e−16
G
CD8 Naive Expression by AML Zone
Anova, p < 2.2e−16
H
Macrophage_Score1
Expression level
CD4 Mem
CD8 Eff
B
Late Erythroid
NK
Platelet/Mega
UnconvT
Monocytes
GMP
pDC
CD8 Naive
cDC
HSC
Early Erythroid
CD4 Treg
CLP
Plasma
CD4 Naive
CD4 Eff
CD8 Mem
AML

Supplemental figure 3. Spatial transcriptomics reveal macrophage proximity to AML cells.

(A) Spatial transcriptomics stratified an AML BM section into 5 zones by distance to AML cells: core_AML, adjacent, proximal, intermediate, and distant.

(B-G) Prediction scores indicate the abundance of AML cells and normal immune populations across zones.

(H) Expression of an independent macrophage gene signature across cell types, confirming highest enrichment in Mono/Mac populations.

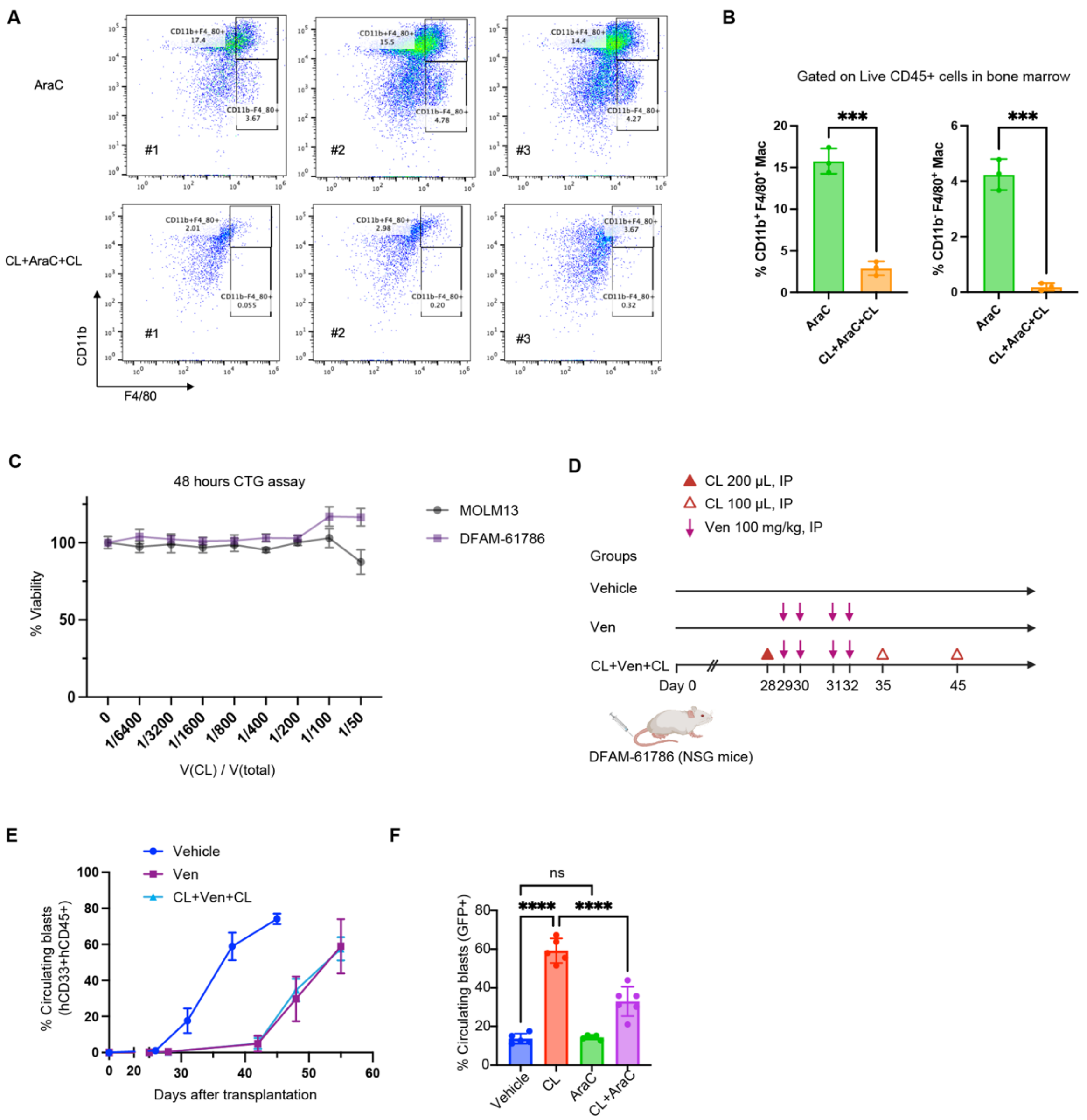

Supplemental figure 4. In vivo macrophage depletion in AML models.

(A) Flow cytometry plots showing gating for macrophages in BM cells from PDX mice treated with AraC or CL+AraC+CL. Cells were gated on live mouse $CD45^{+}$.

(B) Percentage of $CD11b^{+}F4/80^{+}$ macrophages (left panel) and $CD11b^{-}F4/80^{+}$ macrophages (right panel) with or without macrophage depletion in bone marrow of PDX mice.

(C) Viability of MOLM13 and DFAM-61786 AML cells treated with increasing volumes of CL in vitro. CL showed no direct cytotoxicity on AML cells.

(D) Experimental design for Venetoclax (Ven) treatment in DFAM-67186 PDX mice with or without macrophage depletion

(E) Leukemia engraftment curves in PDX mice treated with Ven ± CL. Macrophage deletion did not affect the efficacy of Ven, suggesting that macrophages do not affect resistance to Ven in vivo.

(F) Percentage of blasts in peripheral blood of MLL-AF9 mice with or without macrophage depletion (n = 5-6 mice per group). One-way ANOVA.

*Data are mean ± SD. *, $p < 0.05$; **, $p < 0.01$; ***, $p < 0.001$; ****, $p < 0.0001$.

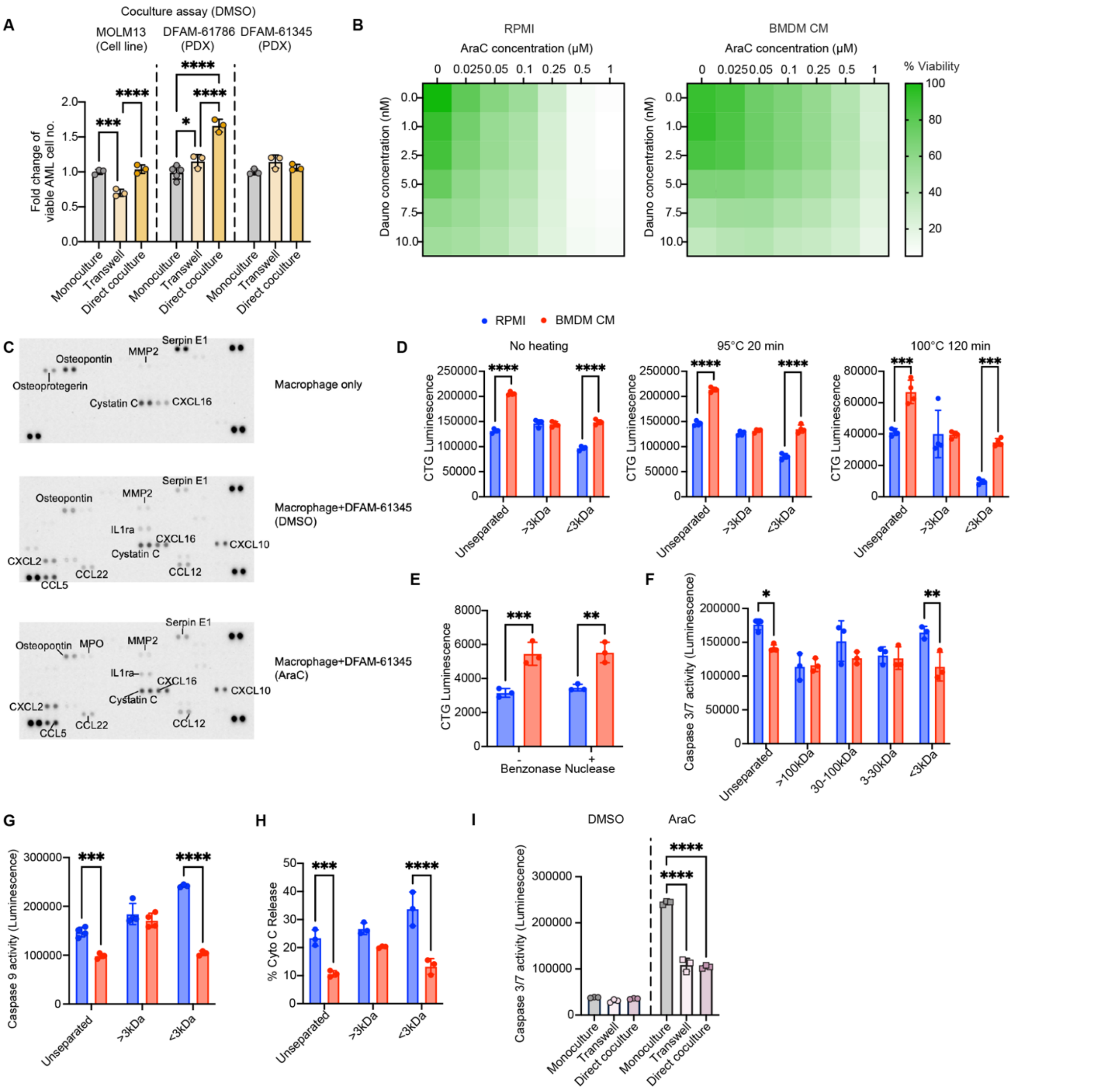

Supplemental figure 5. In vitro assays demonstrate macrophage-derived soluble factors influence AraC efficacy.

(A) Fold change of absolute number of live AML cells after monoculture vs. coculture with macrophages in DMSO control. Fold change is normalized to monoculture treated with DMSO. One-way ANOVA.

(B) Viability of MOLM13 cells treated BMDM CM vs. RPMI control in AraC and Daunorubicin.

(C) Cytokine array of BMDM CM alone, BMDM+DFAM-61345 (DMSO), and BMDM+DFAM-61345 (AraC). BMDM secrete osteopontin which has been previously shown to promote leukemia cell survival.

(D) CTG assay for quantifying viable MOLM13 cells treated with AraC in the presence of BMDM CM or RPMI fractions, with or without heat treatment. Boiling did not abolish the protective activity of CM, indicating that the soluble factors are heat stable.

(E) CTG assay for quantifying viable DFAM-61786 cells treated with AraC in the presence of unseparated BMDM CM or RPMI, with or without Benzonase Nuclease treatment. Benzonase Nuclease did not abolish the protective activity of CM, indicating that the soluble factors are not nucleic acids.

(F-H) Caspase 3/7 activity, Caspase 9 activity, and CytoC release of MOLM13 cells treated with different fractions of BMDM CM or RPMI media. Two-way ANOVA.

(I) Caspase 3/7 activity of monocultured or cocultured MOLM13 cells under DMSO or AraC.

*Data are mean ± SD. *, $p < 0.05$; **, $p < 0.01$; ***, $p < 0.001$; ****, $p < 0.0001$.

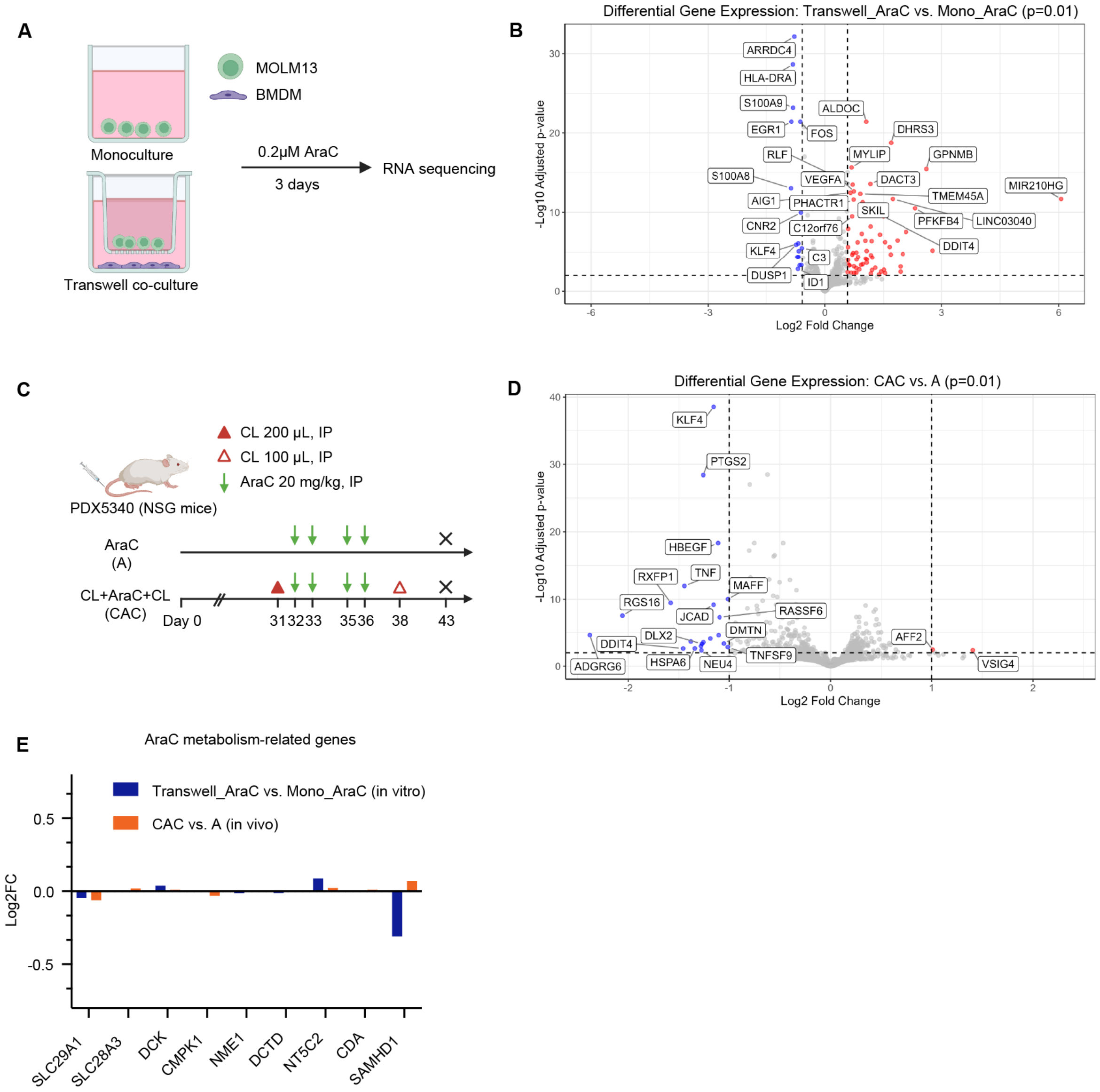

Supplemental figure 6. Transcriptomics showed that macrophages had no influence on the expression of AraC metabolism-related genes in AML cells.

(A) Schematic of RNA-seq experiments on AML cells cultured with or without macrophages.

(B) Volcano plot of differential gene expression in transwell coculture (Transwell_AraC) vs. monoculture (Mono_AraC) (n = 3).

(C) Schematic of RNA-seq on PDX cells separated from mice with or without macrophage depletion.

(D) Volcano plot showing differential expression in PDX cells isolated from CL+AraC+CL (CAC) vs. AraC (A) mice (n = 3).

(E) Log2 FC of AraC metabolism-related genes in both RNA-seq datasets, showing minimal transcriptional changes.

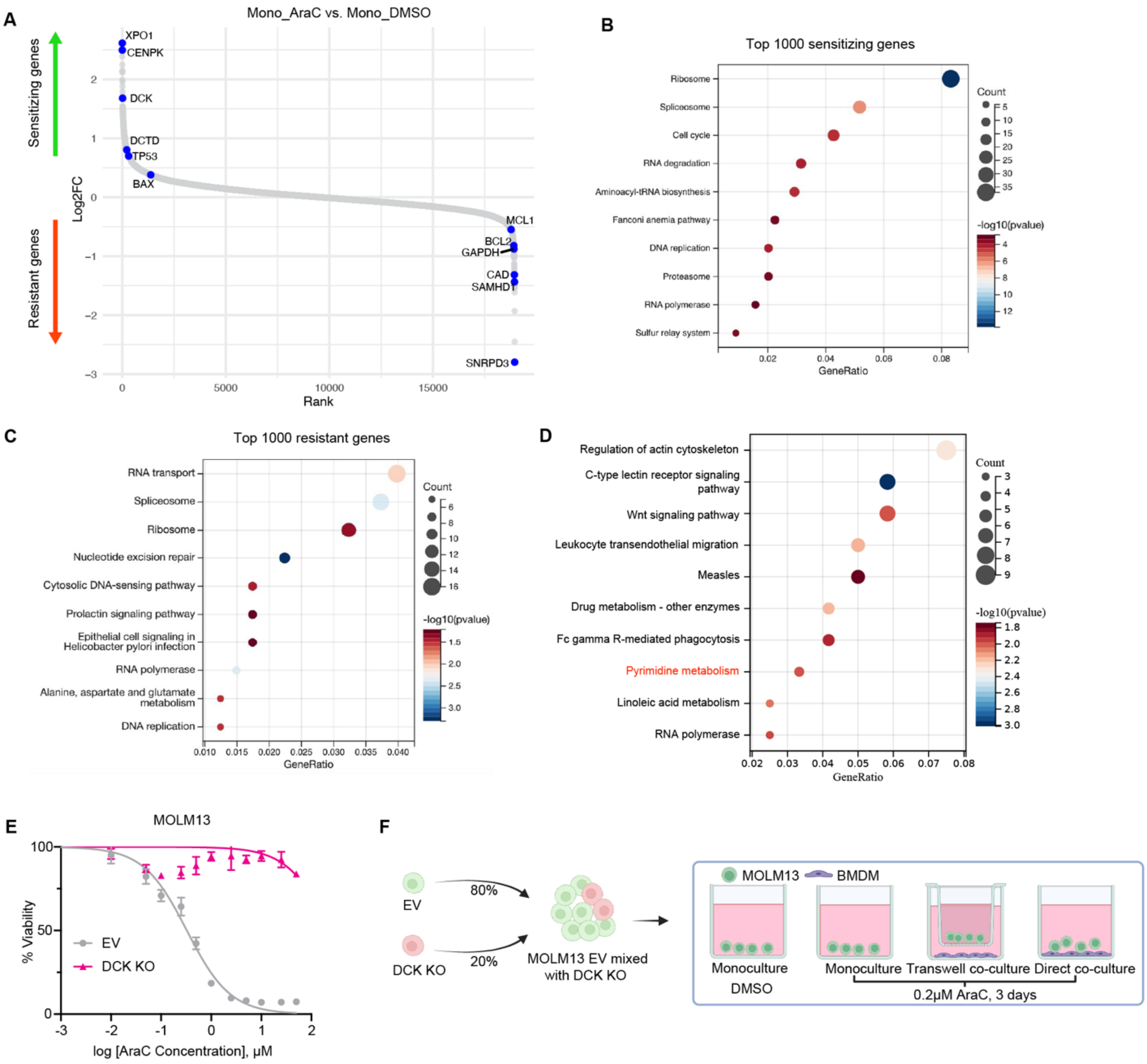

Supplemental figure 7. Genome-wide CRISPR screen identifies DCK as a target of macrophage-derived metabolites.

(A) Log2FC of sgRNA abundance genes comparing Mono_A to Mono_D. The identified AraC-sensitizing and -resistant genes recapitulate previous studies, demonstrating the robustness of the screening.

(B and C) KEGG enrichment analysis of top 1000 sensitizing or resistant genes, recapitulating known tumor-intrinsic AraC resistance pathways.

(D KEGG pathway enrichment of common hits in Figure 3D.

(E) Viability of empty vector (EV) or DCK KO MOLM13 cells under AraC.

(F) Schematic of competition assay validating the CRISPR screen. CellTrace FarRed–labeled DCK KO MOLM13 cells were mixed with empty vector (EV) control cells at a 20% ratio and cultured either alone or with BMDM in the presence of DMSO or AraC.

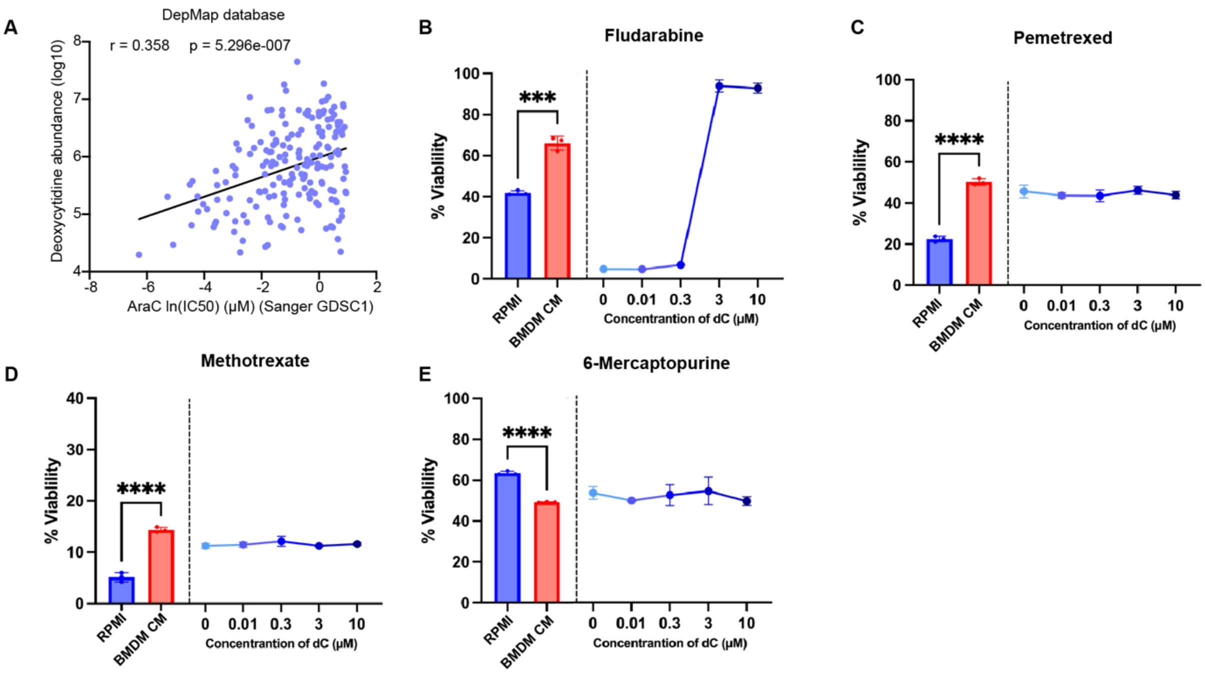

Supplemental figure 8. Macrophages secrete pyrimidine metabolite deoxycytidine (dC) that promotes chemoresistance.

(A) Correlation between dC abundance and AraC IC50 in cancer cell lines (DepMap).
(B-E) Viability of MOLM13 cells treated with RPMI, BMDM CM, or increasing concentrations of dC in the presence of fludarabine (B), pemetrexed (C), methotrexate (D), or 6-mercaptopurine (E).

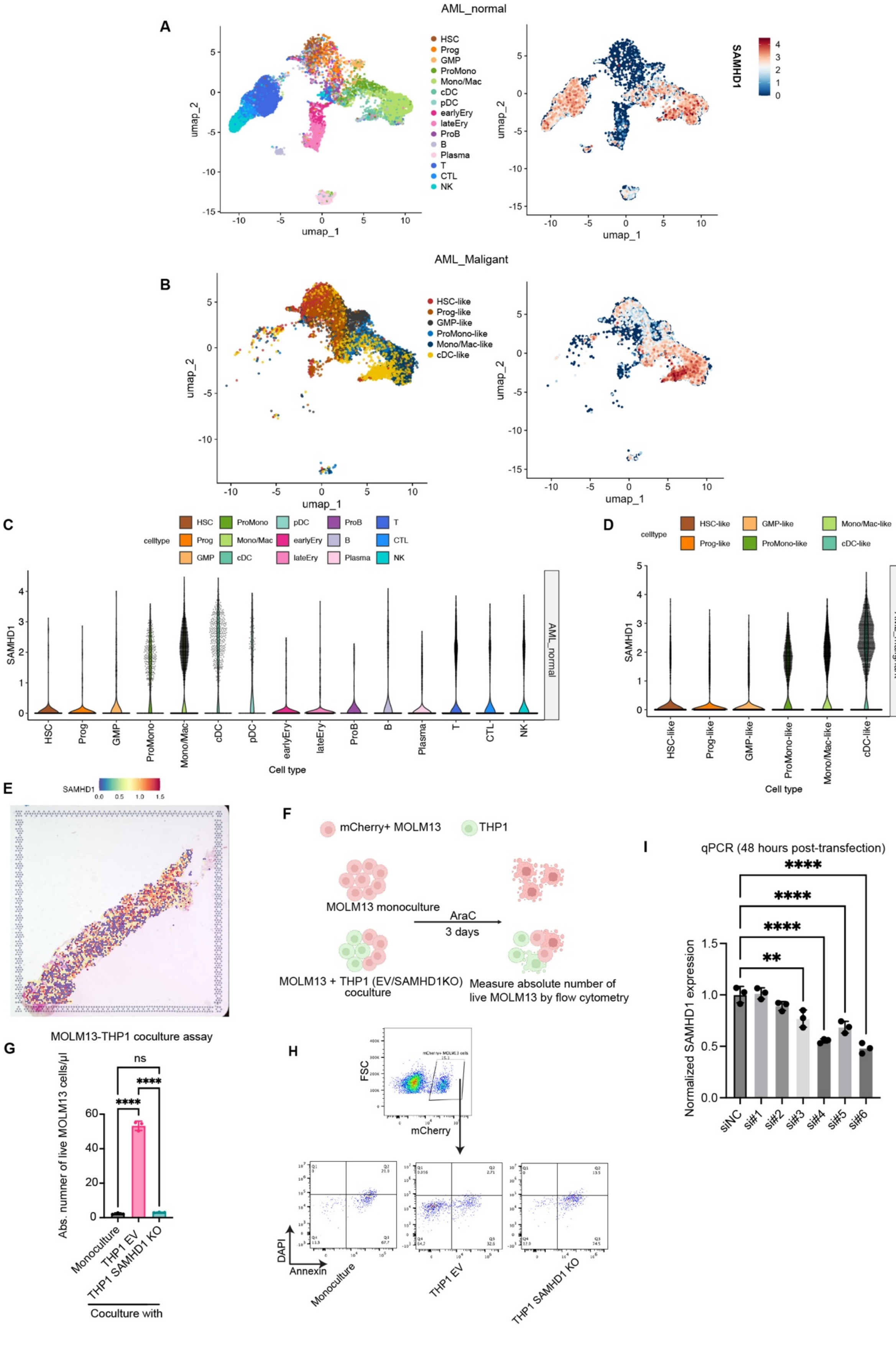

AML_normal
A
HSC
Prog
GMP
ProMono
Mono/Mac
cDC
pDC
earlyEry
lateEry
ProB
B
Plasma
T
CTL
NK
SAMHD1
umap_1
umap_2
AML_Maligant
B
HSC-like
Prog-like
GMP-like
ProMono-like
Mono/Mac-like
cDC-like
C
celltype
Cell type
AML_normal
D
AML_malignant
E
F
mCherry+ MOLM13
THP1
MOLM13 monoculture
AraC
3 days
MOLM13 + THP1 (EV/SAMHD1KO) coculture
Measure absolute number of live MOLM13 by flow cytometry
G
MOLM13-THP1 coculture assay
Abs. numner of live MOLM13 cells/µl
Monoculture
THP1 EV
THP1 SAMHD1 KO
Coculture with
ns
H
FSC
mCherry
DAPI
Annexin
I
qPCR (48 hours post-transfection)
Normalized SAMHD1 expression
siNC
si#1
si#2
si#3
si#4
si#5
si#6

Supplemental figure 9. SAMHD1 drives deoxycytidine (dC) production in macrophages.

(A) UMAP analysis showing the expression of SAMHD1 in normal cells from AML BM (AML_normal).

(B) UMAP analysis showing the expression of SAMHD1 in malignant AML cells (AML_malignant).

(C and D) Quantification of SAMHD1 across different cell types in AML_normal or AML_malignant BM. Highest SAMHD1 expression is observed in DC, followed by Mono/Mac.

(E) Spatial transcriptomic analysis shows the distribution of SAMHD1 expression in AML BM.

(F) Schematic of in vitro coculture with MOLM13 and THP1 EV vs. SAMHD1 KO cells under AraC. mCherry+ MOLM13 cells were cocultured with THP1 EV or SAMHD1 KO cells in the presence of AraC. After three days, the absolute number of viable MOLM13 cells were measured by flow cytometry.

(G) Absolute number of live MOLM13 cells in monoculture vs. coculture with THP1 EV or THP1 SAMHD1 ($n = 3$). One-way ANOVA.

(H) Representative flow cytometry plots showing the gating strategy and MOLM 13 viability after coculture with THP1 cells.

(I) qPCR quantification of SAMHD1 knockdown efficiency in BMDM with six siRNAs targeting mouse SAMHD1. Si#4 and #6 were selected for further experiments since they have the highest KD efficiency. One-way ANOVA.

*Data are mean ± SD. *, $p < 0.05$; **, $p < 0.01$; ***, $p < 0.001$; ****, $p < 0.0001$.

**Key resources table**

| REAGENT or RESOURCE | SOURCE | IDENTIFIER |
|---|---|---|
| Antibodies | | |
| CD45-BV421 | BioLegend | Cat#: 304032; RRID: AB_2561357 |
| CD33-PE | BioLegend | Cat#: 303404; RRID: AB_314348 |
| Zombie Yellow | BioLegend | Cat#: 423104 |
| mCD45-Pacific Blue | BioLegend | Cat#: 103125; RRID: AB_493536 |
| mCD11b-PerCP/Cy5.5 | BioLegend | Cat#: 101228; RRID: AB_893232 |
| F4/80-APC | BioLegend | Cat#: 123116; RRID: AB_893481 |
| Cytochrome C-AF647 | BioLegend | Cat#: 612310; RRID: AB_2565241 |
| DHODH | Cell Signaling Technology | Cat#: 26381 |
| SAMHD1 | Cell Signaling Technology | Cat#: 49158 |
| GAPDH | Cell Signaling Technology | Cat#: 5174; RRID: AB_10622025 |
| Vinculin | Cell Signaling Technology | Cat#: 4650; RRID: AB_10559207 |
| Annexin-AF647 | Thermo Fisher | Cat#: A23204; RRID: AB_2341149 |
| PI | Thermo Fisher | Cat#: 00-6990-50 |
| DAPI | Thermo Fisher | Cat#: 62248 |
| pH2AX-AF647 | Cell Signaling Technology | Cat#: 2577; RRID: AB_2118010 |
| Celltrace FarRed | Thermo Fisher | Cat#: C34564 |
| Biological Samples | | |
| Fresh human leukapheresis blood collars | National University Hospital (NUH), Singapore | |
| Patient blood | National University Hospital (NUH), Singapore<br>MD Anderson Cancer Center (MDACC) | |
| Patient BM cells for scRNA-seq | National University Hospital (NUH), Singapore | |
| Patient BM for spatial transcriptomics | MD Anderson Cancer Center (MDACC) | |

| | | |
|---|---|---|
| Chemicals, peptides, and recombinant proteins | | |
| Cytarabine-in vivo | National University Hospital (NUH), Singapore — Hospital Pharmacy Manufacturer: Pfizer | Presentation: Cytarabine Injection 1 g in 10 mL (sterile) Clear Glass Vial.; Reg. no. (SG): SIN05521P |
| Cytarabine-in vitro | MedChemExpress | Cat#: HY-13605 |
| Cytarabine-d2 | MedChemExpress | Cat#: HY-13605S |
| Clodronate liposomes | LIPOSOMA B.V. | Cat#: C-005 |
| DMSO | Sigma-Aldrich | Cat#: D8418 |
| Lenti-X concentrator | Takara | Cat#: 631232 |
| Polybrene | Sigma-Aldrich | Cat#: H9268 |
| Deoxycytidine | MedChemExpress | Cat#: HY-D0184 |
| Mouse IFN-β1 | Biolegend | Cat#: 581302 |
| Leflunomide | Sigma-Aldrich | Cat#: L5025 |
| Brequniar | TargetMol | Cat#: T8332 |
| Venetoclax | MedChemExpress | Cat#: HY-15531 |
| 4-OH Tamoxifen | Sigma-Aldrich | Cat#: H7904 |
| PEG | Sigma-Aldrich | Cat#: 81172 |
| Tween-80 | Sigma-Aldrich | Cat#: P1754 |
| Mouse M-CSF | Miltenyi Biotec | Cat#: 130-101-703 |
| Human M-CSF | Miltenyi Biotec | Cat#: 130-096-492 |
| Human GM-CSF | Miltenyi Biotec | Cat#: 130-093-865 |
| Human IL-4 | Miltenyi Biotec | Cat#: 130-093-921 |
| Mouse IL-3 | Miltenyi Biotec | Cat#: 130-099-509 |
| Mouse SCF | Miltenyi Biotec | Cat#: 130-101-694 |
| Mouse IL-6 | Miltenyi Biotec | Cat#: 130-096-684 |
| Critical commercial assays | | |
| CellTiter-Glo (CTG) Luminescent Cell Viability | Promega | Cat#: G7572 |
| Caspase 3/7 glo | Promega | Cat#: G8091 |
| Caspase 9 glo | Promega | Cat#: G8211 |
| Mouse cell depletion kit | Miltenyi Biotec | Cat#: 130-104-694 |
| Human CD14 beads | Miltenyi Biotec | Cat#: 130-050-201; RRID: AB_2665482 |
| Proteome Profiler Mouse XL Cytokine Array | R&D Systems (Bio-Techne) | Cat#: ARY006 |
| Experimental models: Cell lines | | |
| MOLM13 | DSMZ | Cat#: ACC-554; RRID: CVCL_2119 |

| | | |
|---|---|---|
| THP1 | ATCC | Cat#: TIB-202; RRID: CVCL_0006 |
| MLL-AF9 | Adam Sperling Lab | |
| DFAM-61786 | | |
| DFAM-61345 | | |
| SB86AML | | |
| SB102AML | | |
| Experimental models: Organisms/strains | | |
| C57BL/6J | InVivos Pte Ltd | JAX Stock No.: 000664; RRID: IMSR_JAX:000664 |
| NSG/JInv (JAX) | InVivos Pte Ltd | AX Stock No.: 005557; RRID: IMSR_JAX:005557 |
| Plasmids | | |
| LentiCRISPR-v2-Puro | Addgene | |
| sgRNA sequences | | |
| DCK | TGTATGAGAAACCTGAACGA | |
| SAMHD1 | TCCATCCCGACTACAAGACA | |
| DHODH | GAGTCTTGAAATCTGGCCCG | |
| Primer sequences | | |
| Mouse SAMHD1 | F: TGC AGA CGA CGA CTT CCA AAA<br>R: TCT CGG AAA CCA CGA TTC TCT AA | |
| Mouse GAPDH | F: TGG CCT TCC GTG TTC CTA C<br>R: GAG TTG CTG TTG AAG TCG CA | |
| siRNA sequences | Sense | Anti-sense |
| siNC | mG*mG*mA mCmGmA fGmGfA fCfGmA mGmCmA mCmU*mU* mC | mG*fA*mA mGmUfG mCmUmC mGmUmC mCfUmC fGmUmC mC*mU*mU |
| mSAMHD1 si#1 | mU*mC*mU mGmUmC fUmGfC fAfGmA mCmGmA mCmG*mA* mC | mG*fU*mC mGmUfC mGmUmC mUmGmC mAfGmA fCmAmG mA*mU*mU |

| | | |
|---|---|---|
| mSAMHD1 si#2 | mG*mA*mG mAmAmU fCmGfU fGfGmU mUmUmC mCmGmA mG*mA*mG | mC*fU*mC mUmCfG mGmAmA mAmCmC mAfCmG fAmUmU mCmUmC* mU*mA |
| mSAMHD1 si#3 | mC*mG*mC mCmGmG fCmUfC fGfUmU mUmCmU mGmC*mC* mC | mG*fG*mG mCmAfG mAmAmA mCmGmA mGfCmC fGmGmC mG*mA*mU |
| mSAMHD1 si#4 | mC*mC*mU mUmAmU fCmAfG fAfAmU mCmAmU mCmGmA mC*mA*mC | mG*fU*mG mUmCfG mAmUmG mAmUmU mCfUmG fAmUmA mAmGmG* mA*mG |
| mSAMHD1 si#5 | mG*mU*mU mCmCmA fGmCfG fAfCmU mUmCmG mCmUmA mU*mA*mU | mA*fU*mA mUmAfG mCmGmA mAmGmU mCfGmC fUmGmG mAmAmC* mU*mG |
| mSAMHD1 si#6 | mG*mU*mG mAmGmC fGmAfG fAfUmA mUmAmC mUmCmU mG*mU*mG | mC*fA*mC mAmGfA mGmUmA mUmAmU mCfUmC fGmCmU mCmAmC* mU*mG |
| Other | | |
| Amicon® Ultra-15 Centrifugal Filter | Merck | Cat#: UFC900324(3KDa), UFC901024(10KDa), UFC903024(30KDa), UFC910024(100KDa) |
| Transwell | Corning | Cat#: 3450 |
| Fetal Bovine Serum （FBS） | Gibco | Cat#: 10082147 |
| Penicillin-streptomycin | Gibco | Cat#: 15140-122 |
| Lipofectamine 3000 | Thermo Fisher | Cat#: L3000015 |
| Lipofectamine RNAiMax | Thermo Fisher | Cat#: 13778150 |
| Lymphosep | Biowest | Cat#: L0560 |
| Glutamax | Gibco | Cat#: 35050-061 |
| MycoStrip® – Mycoplasma Detection Kit | InvivoGen | Cat#: rep-mys-20 |
| RPMI | Thermo Fisher | Cat#: 11875-093 |
| DMEM | Thermo Fisher | Cat#: 11965-092 |
| SFEM II | STEMCELL Technologies | Cat#: 09655 |
| ACK Lysing Buffer | Gibco | Cat#: A1049201 |

| | | |
|---|---|---|
| Benzonase® Nuclease | Merck | Cat#: 70746 |
| Digitonin | Sigma-Aldrich | Cat#: D141 |
| RNeasy kit | QIAGEN | Cat#: 74104 |

**EXPERIMENTAL MODEL AND SUBJECT DETAILS:**

Mice and housing conditions

Mice were purchased from InVivos Pte Ltd. Female NSG mice (6-8 weeks old) were used in PDX tumor graft experiments. Female C57BL/6 mice (6-8 weeks old) were used in MLL-AF9 syngeneic AML graft experiments.

The NUS Institutional Animal Care and Use Committee approved all procedures in accordance with the Guide for the Care and Use of Animals. All mice were housed in pathogen-free facilities, in a 12-hour light/dark cycle in ventilated cages, with chow and water supply ad libitum.

Cell Culture

MOLM13 AML cell line, THP1 human monocytic cell line, RAW264.7 mouse macrophage cell line, HEK293 cell line were purchased from ATCC. Mouse MLL-AF9 AML cells (derived from mouse c-Kit+ cells transduced with Mll-Af9 gene) were kindly provided by Adam Sperling lab. All the cell lines used in tumor studies were confirmed as mycoplasma negative using MycoAlert Mycoplasma Detection kit.

MOLM13 and THP1 cells were cultured and maintained in complete RPMI media (10% fetal bovine serum (FBS), 100 units/ml penicillin-streptomycin). MLL-AF9 cells were maintained in complete RPMI media supplemented with 10 ng/ml mouse IL-3. RAW264.7 and HEK293 cell lines were cultured in complete DMEM media (10% FBS, 100 units/ml penicillin-streptomycin, glutamax). PDX cells were maintained in complete RPMI media for a short term ($\leq$ 3 days). Primary AML cells were maintained in 25% SFEM II media (with 1X StemSpan™ CD34+ Expansion Supplement) + 75% complete RPMI media for a short period ($\leq$ 3 days).

BMDM cells were differentiated from the bone marrow cells of NSG or C57BL/6 mice. Briefly, bone marrow cells were harvested from tibia and femur of mice using mortar and pester, followed by RBC lysis using LCK lysis buffer [1]. The bone marrow cells were then resuspended in complete RPMI media supplemented with 50 ng/ml mouse M-CSF at 8x10^5/ml and then seeded in well plates at a suitable volume. Half of the media were replaced on day 4. On day 6, all media were replaced. On day 7, the BMDM are ready to use.

hMDM were differentiated from human peripheral blood CD14+ monocytes. PBMC from healthy donors were separated from blood using Lymphosep (Biowest). CD14+ monocytes were separated from PBMC using CD14 beads (Miltenyi) following the product manual. The CD14+ monocytes were then resuspended in complete DMEM media supplemented with 50ng/ml human M-CSF at a density of 5x10^5/ml and then seeded in suitable containers at a proper volume. Half of the media were replaced on day 4. On day 6, all media were replaced. On day 7, the hMDM are ready to use.

#### METHOD DETAILS

Patient-derived xenograft (PDX) AML model

6x10^5 DFAM-61786 cells/mouse were resuspended in 200 ul of PBS and injected intravenously into the tail vein of NSG mice. Mice were randomized using Excel random number generator before the treatment. Tumor growth was monitored every week by bleeding from the submandibular vein [1]. Mice were euthanized when the percentage of tumor cells reached 70% in peripheral blood or when they had symptoms and reached the humane endpoint.

MLL-AF9 syngeneic AM model

5x10^5 GFP+ MLL-AF9 cells/mouse were resuspended in 200 ul of PBS and intravenously injected into the tail vein of C57BL/6 mice. Mice were randomized using Excel random number generator before the treatment. Tumor growth was monitored every week by bleeding from the submandibular vein. Mice were euthanized when the percentage of tumor cells reached 70% in peripheral blood, or they had symptoms and reached humane endpoint.

Macrophage depletion

200 ul of Clodronate Liposomes (Liposoma) was intraperitoneally injected as induction depletion. 100 ul of CL was intraperitoneally injected every 7 days as maintenance. Macrophage depletion was validated by flow cytometry analysis of bone marrow cells.

Flow cytometry analysis of mouse tumor burden

Tumor burden was monitored every week using the peripheral blood collected from submandibular vein. To lyse red blood cells, 13 µl of blood was added into 300 µl of ACK lysis buffer (Gibco) and incubated at room temperature for 9 minutes, followed by adding 2 ml of PBS. The white blood cells were then pelleted down by centrifugation at 500 g for 5 minutes. Cell pellet was then resuspended in 100 µl PBS before staining. For the PDX model, cells were stained with 1% Zombie Yellow (Biolegend), 1% human CD45-BV421 (Biolegend), and 0.5% human CD33-PE (Biolegend) for 20 minutes. For MLL-AF9 syngeneic model, cells were stained with 1% Zombie Yellow for 20 minutes. After staining, samples were diluted with 200 µl of PBS and measured by flow cytometry. For PDX model, tumor cells were identified as Zombie Yellow-hCD33+hCD45+. MLL-AF9 cells were identified as Zombie Yellow-GFP+.

To analyze tumor burden at the endpoint, single cell suspension of bone marrow and spleen was collected by mortar and pestle or by mincing and through a mesh strainer, respectively [1]. RBCs were then lysed using ACK lysis buffer. After the lysis, cells were then pelleted down and resuspended in 1x Benzonase® Nuclease (Merck) PBS buffer and digested at room temperature for 15 – 20 minutes followed by filtration through mesh strainer to remove cell clumps. Around 3-5 millions of cells were then used for flow cytometry. PDX samples were stained with 1% Zombie Yellow, 0.5% hCD33-PE, and 1% hCD45-BV421. MLL-AF9 cells were stained with 1% Zombie Yellow PBS buffer. After staining at 4 degree for 30 minutes, cells were washed once with 900 µl of cold PBS and acquired by flow cytometry.

Western blot

Cells were lysed in NP-40 lysis buffer (150 mM NaCl, 1% NP-40, 50 mM Tris-Cl pH 8.0; Cell Signaling Technology) supplemented with protease inhibitor for 45 minutes at 4°C according to the manufacturer's protocol. Protein concentration was determined using Pierce BCA Protein Assay Kit (ThermoFisher). Proteins containing 10% β-mercaptoethanol (Sigma-Aldrich) were boiled at 95°C for 10 minutes, separated by SDS-PAGE, and transferred onto nitrocellulose membranes. Membranes were blocked for 1 hour with 5% milk (Bio-Rad) dissolved in TBST (w/v) and incubated overnight in primary antibodies diluted in 5% milk. List of antibodies used are listed in key source table.

##### qPCR

Total RNA was extracted using RNeasy mini kit (QIAGEN) and treated with DNase I (QIAGEN). For RT-qPCR, a total of 1 μg of purified RNA was reverse-transcribed into cDNA using GoScript Reverse Transcriptase (Promega). qPCR was performed using Power SYBR Green PCR Master Mix (Applied Biosystems) on the CFX96 Touch Real-Time PCR Detection System (BioRad). mRNA expression levels were evaluated using the ΔΔ Ct method. Primer sequences are listed in key source table.

##### Macrophage AML coculture assays

For BMDM-AML coculture assays, bone marrow cells from NSG mice were differentiated into BMDM as described above. Mature BMDM were then dissociated from well plates using 2mM EDTA in PBS. 1 ml of BMDM in complete RPMI media at a density of 3x10^5/ml were seeded in wells of 12 well plate. 0.5 ml of AML cells in complete RPMI media at a density of 6x10^5/ml were then added to BMDM either as direct coculture or in the upper chamer of a transwell insert. For monoculture, tumor cells were added to 1 ml complete RPMI media. AraC or DMSO was prediluted in both BMDM or tumor cell suspension at a suitable concentration. After 3 days of coculture, AML cells, which are suspension cells, were harvested into falcon tubes, pelleted down, and supernatant was carefully discarded without disturbing the cell pellet. After the staining with Annexin-Alexa Fluor 647 (Invitrogen), DAPI (Invitrogen), and CD33-PE (Biolegend), cells were diluted with 200 μl of PBS. Absolute number of Live CD33+ AML cells were measured by CytoFLEX flow cytometer (Beckman Coulter).

For hMDM-AML coculture assays, hMDM were differentiated from CD14+ monocytes in 96 well plates described above. On day 7, supernatant of macrophages was removed, and adherent macrophages were washed with complete RPMI media, followed by adding 100 μl of complete RPMI media supplemented with DMSO, AraC, or Ven. 6x10^4 AML cells in 100 μl of complete RPMI media were then added to the macrophages. For AML monoculture control, the same number of AML cells were added to 100 μl media supplemented with drugs. After 3 days of coculture, remaining suspension cells were harvested, pelleted down, and stained with Annexin-Alexa Fluor 647, DAPI, and CD33-PE. Cells were then diluted with 200 μl of PBS. Absolute number of Live CD33+ AML cells were measured by CytoFLEX flow cytometer (Beckman Coulter).

#### Generating CM

BMDM or hMDM were differentiated as described above for 7 days in 12 well plates. On day 7, old media were replaced with 2 ml/well of fresh complete RPMI or DMEM media for BMDM and hMDM, respectively. After three days, coarse CM supernatant was harvested. For cell lines that grow exponentially, including MOLM13, THP1, and RAW264.7, 3x10^5/ml cells were seeded in their respective culture media in well plates. For PDX cells, 6x10^5/ml cells were seeded in complete RPMI media in well plates. CM were harvested after three days of culture.

The coarse CM were first centrifuged at 1000 g and then filtered through 0.4 μm syringe filter to remove cellular debris. CM were aliquoted and stored in -80°C for further assays.

#### Fractioning experiments of CM

To fraction CM and complete RPMI media, the samples were filtered through a series of Amicon® Ultra-15 Centrifugal Filter Units from large to small molecular weight. For example, 10 ml of samples were first filtered through 100kDA filter and concentrated into 200 μl. The concentrated >100kDa fraction was reconstituted with complete RPMI media into the initial volume, namely 10 ml here. The filter through (<100kDa) fraction was then filtered with a second Amicon filter (30kDa). The process was repeated until samples had passed through a 3kDa filter. All centrifugation was done at 3000 g, 4°C. All fractions were aliquoted and stored in -80.

#### CTG assay and Caspase glo assays

Cell viability was assessed using the CellTiter-Glo (CTG) Luminescent Cell Viability Assay (Promega), which quantifies ATP levels as a marker of metabolically active cells. On day 0, AML cells were seeded at 20,000/ml in 200 μL of culture medium in 96-well plates or in 60 μL of culture media in 384-well plates. Cells were treated with various concentrations of test compounds and incubated for 72 hours at 37°C with 5% $CO_2$. To test the CM of macrophages, cells were treated with 50% of CM and various concentrations of compounds.

After three days, 15 μL of CTG reagent was added directly to each well of a 96-well plate, or 5 μl of reagent as added to each well of a 384-well plate, followed by gentle shaking on an orbital shaker at 200 rpm for 10 minutes at room temperature in the dark to ensure complete cell lysis and mixture. Luminescence was measured using a Hidex Sense multi-mode plate reader.

Caspase activity was measured using the Caspase-Glo® 3/7, 8, and 9 Assay Kits (Promega), which provide a luminescent readout proportional to caspase activation. Cells were seeded at 6,000 cells per well in 30 μL of culture medium with or without 50% BMDM CM in 384-well plate and treated with AraC for 4 hours. At the endpoint, 30 μL of the corresponding Caspase-Glo® reagent was added directly to each well, and the plate was gently mixed on an orbital shaker at 200 rpm for 1 hour at room temperature in the dark. Luminescence was measured using a Hidex Sense multi-mode plate reader.

Flow cytometry staining

For surface staining, around 1x10^6 cells were harvested and pelleted down by centrifugation and then resuspended in 100 μL PBS containing the fluorophore-labeled antibodies and incubated with at room temperature in the dark for 25 minutes, cells were washed with cold PBS and resuspended in 200 μL PBS for FACS acquisition.

For p-H2AX intracellular staining, cells were first treated with AraC with or without CM for 4 hours and then harvested and pelleted down at 400 × g for 5 minutes. The pellet was then resuspended in 100 μl Cytofix/Cytoperm solution (BD) and fixed at 4°C for 20 minutes. After fixation, cells were washed twice with 500 μl 1X Perm/Wash buffer (BD) and then incubated in 100 μl 1X Perm/Wash buffer with 2 μl anti-p-H2AX-Alexa Fluor 647 antibody (CST) at room temperature in the dark for 25 minutes. After the staining, the cells were then washed twice with 500 μl 1X Perm/Wash buffer and resuspended with 100 μl 1 μg/ml DAPI in PBS and incubated at room temperature for 20 minutes in the dark before FACS analysis.

CytoC release assay

The CytoC release assay was adapted from BH3 profiling as described in previous literatures (cite Shruti's paper). Briefly, cells were treated with AraC with or without BMDM CM for 18-20 hours. Cells were then pelleted down and permeabilized in MEB buffer (150mM mannitol, 10mM HEPES-KOH pH 7.5, 50mM KCl, 0.02mM EGTA, 0.02mM EDTA, 0.1% BSA and 5mM Succinate) with digitonin (0.002%) for 60 minutes. After 60 minutes peptide exposure at room temperature, cells were fixed using 4% formaldehyde for 15 minutes, followed by neutralization for 10 minutes using N2 buffer (1.7M Tris, 1.25M Glycine pH 9.1). Sensitivity to BH3 peptides were measured as cytochrome c loss using anti-cytochrome c Alexafluor 647 antibody (Clone 6H2.B4, Biolegend) in staining buffer (10% BSA, 2% Tween20, PBS) via FACS by a gating strategy, in which a gate was drawn around the DMSO-negative control to depict 100% cytochrome c retention.

#### Metabolomics

For sample preparation, 100 μL of the sample was mixed with 300 μL of pre-cooled (−80°C) 80% methanol, vortexed for 1 minute, incubated at -20°C for 2 hours and centrifuged at 16,000 g for 20 minutes at 4°C. The supernatant was carefully transferred and further filtered through an 0.22 μm filter before analysis. Extraction solution was dried using SpeedVac. 60% acetonitrile was added to the tube for reconstitution following by overtaxing for 30 sec. Samples solution was then centrifuged for 30 min @ 20,000g, 4 °C. Supernatant was collected for LCMS analysis.

Samples were analyzed by High-Performance Liquid Chromatography and High-Resolution Mass Spectrometry and Tandem Mass Spectrometry (HPLC-MS/MS) as described in a previous literature [2]. Specifically, system consisted of a Thermo Q-Exactive in line with an electrospray source and an Ultimate3000 (Thermo) series HPLC consisting of a binary pump, degasser, and auto-sampler outfitted with a Xbridge Amide column (Waters; dimensions of 3.0 mm × 100 mm and a 3.5 μm particle size). The mobile phase A contained 95% (vol/vol) water, 5% (vol/vol) acetonitrile, 10 mM ammonium hydroxide, 10 mM ammonium acetate, pH = 9.0; B was 100% Acetonitrile. The gradient was as following: 0 min, 15% A; 2.5 min, 64% A; 12.4 min, 40% A; 12.5 min, 30% A; 12.5-14 min, 30% A; 14-21 min, 15% A with a flow rate of 150 μL/min. The capillary of the ESI source was set to 275 °C, with sheath gas at 35 arbitrary units, auxiliary gas at 5 arbitrary units and the spray voltage at 4.0 kV. In positive/negative polarity switching mode, an m/z scan range from 60 to 900 was chosen and MS1 data was collected at a resolution of 70,000. The automatic gain control (AGC) target was set at 1 × 106 and the maximum injection time was 200 ms. The targeted ions were subsequently fragmented, using the higher energy collisional dissociation (HCD) cell set to 30% normalized collision energy in MS2 at a resolution power of 17,500. Besides matching m/z, target metabolites are identified by matching either retention time with analytical standards and/or MS2 fragmentation pattern. Data acquisition and analysis were carried out by Xcalibur 4.1 software and Tracefinder 4.1 software, respectively (both from Thermo Fisher Scientific).

#### Quantification of dC in CM and AraC-d2 in gDNA by HPLC

For CM sample preparation, 100 μL of the sample was mixed with 300 μL of pre-cooled (−80°C) methanol, vortexed for 5 minutes, and centrifuged at 16,000 g for 20 minutes at

4°C. The supernatant was carefully transferred and further filtered through a 0.22 µm HPLC filter before analysis.

For measurement of intracellular AraC-d2 abundance, the cells were first treated with 0.5 µM AraC-d2 (MCE) and BMDM CM for 22 hours. At the end point, the cells were pelleted down. Genomic DNA was isolated following the manual of genomic DNA extraction kit (NEB). 1 µg purified genomic DNA was then digested into single nucleotides using the NEB Nucleoside Digestion Mix (NEB #M0649)). The digested sample was then mixed with 3 volumes of pre-cooled (-80°C) methanol, filtered through a 0.22 µM HPLC filter before analysis.

The chromatographic separation was performed using a triple quadropole 3500 system from Sciex (Framingham, MA, USA) couples with an Agilent HPLC 1290 system (Santa Clara, CA, USA). The chromatographic separation was performed on C18 100 mm x 2.1 mm, 1.8 µm, column (Waters). Mobile phase A consisted of ammonium formiate (10mM) in water containing 0.1% (v/v) formic acid and mobile phase B consisted of acetonitrile.

Cytokine arrays

Cytokines in the CM of BMDM CM were measured by Proteome Profiler Mouse XL Cytokine Array (R&D) following the product's manual.

CRISPR screen

A genome-wide CRISPR knockout screen was conducted using the Brunello sgRNA library, which targets approximately 19,000 human genes with an average of four sgRNAs per gene (cite). MOLM13 cells were transduced with the lentiviral library, followed by puromycin selection for 7 days to establish a stable knockout population. The screen was performed under experimental and control conditions while maintaining a minimum 300-fold representation of the library at all time points.

The MOLM13 cells carrying the sgRNA library were cultured alone or together with BMDM in 6 well plates with or without transwell insert in the presence of 0.2µM AraC. Suspension cells were harvested after 3 days. Cell pellets were frozen at -40°C before genomic DNA extraction.

Genomic DNA was extracted using phenol:chloroform:isoamyl alcohol (25:24:1) and MaXtract High Density (QIAGEN). The sgRNA sequences were amplified via PCR, purified with AMPure XP beads, and quantified using a Qubit 4 Fluorometer before sequencing on an Illumina NovaSeq 6000 platform. Sequencing reads were processed by removing adapters and aligning to the reference library, and sgRNA enrichment or depletion was analyzed using MAGeCK to identify genes associated with cell viability or treatment.

Collection of mouse BM aspirate

Mouse BM aspirates were obtained from tibia and femur bones using a centrifugation-based method as shown in figure 5A. Briefly, bones were harvested, cleaned in PBS, and cut into two segments at the mid-shaft. Each bone fragment was placed with the cut end down into a perforated 0.2 mL PCR tube nested within a 1.5 mL microcentrifuge tube containing 30 µl of PBS. Samples were centrifuged at 5,000 × g for 30 s at 4°C to expel BM. Residual marrow was dislodged if necessary and centrifuged again. The aspirate was clarified by centrifugation at 500 g for 5 min, followed by 16,000 × g for 10 min at 4°C, and the supernatant was collected for immediate use or storage at –80°C.

CRIPSR KO in cell lines

The protocol of gene CRISPR KO in cell lines have been described in a previous paper [3]. Briefly, sgRNAs were synthesized and cloned into LentiCRISPR-v2-Puro vector. The LentiCRISPR-sgRNA plasmid was then co-transfected with pSPAX2 and pVSV plasmids into HEK293 cells to package LentiCRISPR-sgRNA lentivirus. The coarse lentivirus supernatant was harvested, filtered through 0.45µM filter, and then concentrated using Lenti-X concentrator (Takara). Cell lines were infected with LentiCRISPR-sgRNA lentivirus for 2 days before selection in 2 µg/ml Puromycin (Sigma). After 6 days of selection with Puromycin, gene knock out was validated with WB or qPCR.

Genetic deletion of Dhodh in BMDM by 4-OH Tamoxifen

BMDM from WT or tamoxifen-inducible Dhodh knockout mice ($Dhodh^{fl/fl}$; Cre-ERT2) were differentiated as described above but in the presence of 4µM 4-OH Tamoxifen (MCE) to induce Dhodh KO. On day 7, cells were washed with PBS once and cultured in complete RPMI media for 2 days to generate CM.

SAMHD1 knockdown in BMDM by siRNA

We incorporated phosphorothioates on the last 2 nucleotides of each end of the sense and anti-sense strands. We incorporated 2’-O-methyl and 2’-Fluoro ribose modifications based on patterns known to increase metabolic stability [4]. BMDMs were differentiated from mouse bone marrow cells as described above in RPMI complete media supplemented with mouse M-CSF (Miltenyi). On day 5 or 6 of differentiation, BMDMs were transfected with siRNAs using Lipofectamine RNAiMax (Invitrogen) following the product manual.

#### Gene signature scores

To accurately determine the relative contribution of the immune population in the cohort, we selected for primary AML specimens from bone marrow aspirates with less than 30% blasts population (n = 57). We normalized bulk RNA-seq raw counts using variance stabilizing transformation (VST). Immune gene signatures were calculated using single-sample gene set expression analysis (ssGSEA), using LM22 gene sets from Chen et al., 2018 and gene sets from Ruben Bill et al., 2023.

#### Conditional Inference Tree Analysis

Gene signature scores were analyzed using a conditional inference tree (ctree function, partykit R package) to identify significant predictors of overall survival. The tree was limited to a single split (max depth = 1, minbucket = 20) with a significance threshold of $p < 0.05$. For each significant predictor, the derived split point was used to classify patients into high- and low-score groups, and overall survival differences were visualized using Kaplan – Meier curves.

#### Single cell RNA-seq

We analyzed single-cell RNA sequencing (scRNA-seq) from 12 samples obtained before and after clardribine and low-dose cytarabine treatment from 6 AML patients (24,820 cells total) generated at MD Anderson. Single-cell libraries were prepared using the Chromium Single Cell 3' Reagent Kits v3 (10x Genomics) and sequenced on an Illumina platform. Data processing and quality control were carried out using the Cell Ranger software. High-quality cells with a minimum of 500 genes and less than 10% mitochondrial RNA content were retained for preprocessing. Data normalization, scaling, and clustering (Leiden algorithm) were carried out using the Scanpy package (v1.7.2). Cell type annotation was performed using the CellTypeST tool, based on known marker genes.

Published dataset

Healthy bone marrow scRNA-seq data on were obtained from van Galen et al., 2019 (GSE116256) were used following preprocessing and analysis.10 The dataset included scRNA-seq count data of 38,410 cells from 35 acute myeloid leukemia (AML) samples derived from 16 patients, as well as 5 healthy bone marrow controls. Data preprocessing was performed using Seurat with default parameters for normalization, scaling, and clustering before visualizing using UMAP. Harmony batch removal was performed which accounted for patient-dependent batch effect. To correct for patient-specific batch effects, Harmony was applied for batch correction. Cell type annotations were retained as defined in the original study. Only SAMHD1 expressions on healthy cells were displayed.

RNA-seq

Total RNA of cells was extracted using RNeasy kit (Qiagen). For library construction, mRNA was captured by magnetic beads with Oligo (DT). The first strand of cDNA was synthesized in the M-MuLV reverse transcriptase system using fragmented mRNA as a template and random oligonucleotides as primers. RNaseH degraded the RNA strand, and the second cDNA strand was synthesized by dNTPs in the DNA polymerase I system. The double-stranded cDNA was purified and subjected to end repair, the dA-tail was added, and the sequencing joint was connected. About 200bp cDNA was purified with AMPure XP beads, amplified by PCR, and purified again with AMPure XP beads to obtain the final sequencing library. After the construction of the library, the quality of the library was examined by Kapa qPCR quantification and Agilent 4200 TapeStation analysis. After passing the library inspection, different libraries are pooled according to the requirements for effective concentration and targeted downstream data volume and then sequenced by Illumina NovaSeqX Plus 25B PE150.

Raw RNA-seq data were processed utilising default parameters for TrimGalore 0.6.6 to perform quality control and trimming of low-quality reads and adapters. Alignment of reads was executed using STAR 2.7.5b with parameters --outSAMtype BAM SortedByCoordinate --outSAMunmapped Within --outSAMattributes NH HI NM MD AS. For the reference genome, the GRCh38 primary assembly provided by GENCODE was employed. Gene assignments were based on the GENCODE version 44 primary assembly annotation. Read counts were generated directly from the aligned BAM files using STAR's --quantMode GeneCounts option. RNA-seq data from PDX models were aligned to the mouse reference genome and processed with XenofilterR to remove mouse-derived reads prior to generating read counts with featureCounts using default parameters for paired-end data.

Genes with gene types annotated as pseudogene or artifact and gene id with "PAR_Y", and those for which counts were <1 CPM or exhibited no variance were excluded. Differential gene expression analysis between selected groups was performed using DESeq2.

Quantification and statistical analysis

Calculation of statistical significance was conducted using two-tailed t-tests unless otherwise specified. Differences in cumulative survival probability was determined by Kaplan-Meier Analysis using Log-rank test. For all figures, *p<0.05, **p<0.01, ***p<0.001, ****p<0.0001
